# Glioblastoma-infiltrating CD8^+^ T cells are predominantly a clonally expanded *GZMK*^+^ effector population

**DOI:** 10.1101/2023.08.25.554678

**Authors:** Anthony Z. Wang, Bryce L. Mashimo, Maximilian O. Schaettler, Ngima D. Sherpa, Lydia A. Leavitt, Alexandra J. Livingstone, Saad M. Khan, Mao Li, Markus Anzaldua-Campos, Joseph D. Bradley, Eric C. Leuthardt, Albert H. Kim, Joshua L. Dowling, Michael R. Chicoine, Pamela S. Jones, Bryan D. Choi, Daniel P. Cahill, Bob S. Carter, Allegra A. Petti, Tanner M. Johanns, Gavin P. Dunn

## Abstract

Recent clinical trials have highlighted the limited efficacy of T cell-based immunotherapy in patients with glioblastoma (GBM). To better understand the characteristics of tumor-infiltrating lymphocytes (TIL) in GBM, we performed cellular indexing of transcriptomes and epitopes by sequencing (CITE-seq) and single-cell RNA sequencing (scRNA-seq) with paired V(D)J sequencing, respectively, on TIL from two cohorts of patients totaling 15 patients with high grade glioma, including GBM or astrocytoma, IDH mutant, grade 4 (G4A). Analysis of the CD8^+^ TIL landscape reveals an enrichment of clonally expanded *GZMK*^+^ effector T cells in the tumor compared to matched blood, which was validated at the protein level. Furthermore, integration with other cancer types highlights the lack of a canonically exhausted CD8^+^ T cell population in GBM TIL. These data suggest that *GZMK*^+^ effector T cells represent an important T cell subset within the GBM microenvironment and which may harbor potential therapeutic implications.

**Significance:** In order to understand the limited efficacy of immune checkpoint blockade in GBM, we endeavor to understand the TIL landscape through a multi-omics approach. In this study, by highlighting the enrichment of *GZMK*^+^ effector T cells and lack of exhausted T cells, we provide a new potential mechanism of resistance to immunotherapy in GBM.

## Introduction

Glioblastoma (GBM) is a highly aggressive brain cancer which is the most common primary malignancy of the central nervous system (CNS) in adults (1). Because the prognosis for patients diagnosed with GBM remains poor, there is a critical need for improved treatments. Given the increasing role and success of immunotherapies in many different cancer types, there has been tremendous enthusiasm to explore immune-based approaches to treat GBM. Despite promising results in early phase studies, there currently are no FDA approved immunotherapies for GBM due to lack of efficacy demonstrated in multiple randomized phase 3 clinical trials (2–4). For example, anti-PD-1 treatment, either as monotherapy or in combination, did not improve overall survival in newly diagnosed and recurrent GBM. Similarly, EGFRvIII-specific peptide vaccination also did not significantly improve overall patient survival (5).

Ongoing work is directed at trying to understand the barriers to treatments that are designed to license, or activate, endogenous T cell activity against brain tumor cells. Indeed, it is possible that both tumor intrinsic as well as microenvironmental characteristics of GBM tumors may prevent the immune system from being unleashed in a therapeutically effective manner. Recent work has revealed the significant somatic variant, neoantigen, and cellular intratumoral heterogeneity of GBM, underscoring the difficulty in targeting clonal antigens and specific cell types with disparate transcriptional and functional profiles (6–10). Furthermore, we speculate that this immunologically unfavorable environment represents a particularly severe example of cancer immunoediting in humans (11,12). A myriad of immune deficits have been cataloged including intrinsic immune suppression (13–15) and overexpression of several checkpoint molecules, such as PD-L1 (7,16). At the level of the tumor extrinsic microenvironment, the use of single cell analysis has demonstrated that GBM is largely immunosuppressive in both murine and human settings (17–20).

GBM is often described as a “non-inflamed” tumor because of both the quantitative lack of a robust *de novo* T cell infiltrate as well as the qualitative dysfunction of T cells that are present within the tumor. Indeed, the T cells in GBM are typically considered to be “exhausted” due to expression of surface receptors such as PD-1, LAG-3, and TIM-3 in human samples and functional hypoactivity in mouse models (21–23). One recent study has investigated the sex-biased difference of T cell exhaustion in patient GBM samples, highlighting the difference in frequency of progenitor exhausted T cells and TOX expression between tumor infiltrating T cells derived from male and female patients (24). However, prior studies describing T cell hypofunctionality in human GBM samples have not identified significant transcriptional exhaustion signatures. For example, Mathewson *et al.* (17) characterized a subset of T cells that expressed *KLRB1* (encoding CD161). While inactivation of CD161 enhanced anti-tumor T cell function, there was a noticeable lack of RNA expression of several canonical exhaustion surface markers, such as *TIGIT, LAG3, HAVCR2,* and *CTLA4*, represented by an inhibitory signature. Additionally, a separate study by Abdelfattah *et al.* (25) performed single cell RNA sequencing (scRNA-seq) on several glioma types, including GBM, and similarly showed low expression of several exhaustion markers, such as *LAG3, TIGIT,* and *CTLA4*. Finally, a recent study by Ravi *et al.* (20) implicating the role of myeloid IL-10 as a driver of T cell exhaustion in GBM observed a very low frequency of T cells that harbored canonical markers of exhaustion, such as *LAG3, HAVCR2, CTLA4,* and *PDCD1.* In the aforementioned sex-biased T cell exhaustion study, the total frequencies of both progenitor and terminally exhausted T cells represented a minority of the overall total T cell landscape. Thus, these studies suggest that conventional T cell exhaustion may not be a predominant phenotype of GBM tumor-infiltrating lymphocytes (TIL) and highlight the need for further work to characterize the T cell populations found within brain cancers on a molecular level.

To better understand the T cell compartment within GBM, we sought to characterize T cell phenotypes and cell states using a combined multi-omics approach. To this end, we integrated single cell RNA sequencing (scRNA-seq) with paired antibody capture and single cell V(D)J sequencing on T cells isolated from a cohort of patients diagnosed with malignant gliomas comprised mostly of GBM but several cases of astrocytoma, IDH mutant, grade 4 (G4A) as well. We show that the canonical T cell exhaustion signature was weakly expressed in HGG TIL, which further supports prior studies (17,26). Notably, we observe that HGG TIL are enriched for CD8^+^ *GZMK*^+^ T cells that do not highly express classic cytotoxic markers such as *PRF1, GNLY,* and *GZMB*. Moreover, CD8^+^ *GZMK^+^* T cells can be further subdivided with respect to *NR4A2* expression. Using TCR clonotype data, we find that *GZMK*^+^ T cells are selectively expanded in TIL but not in matched PBMC. By integrating T cell expression data from a broad range of cancer types, including an independent GBM cohort, we gain further validation supporting the lack of exhausted T cells in HGG as well as the relative increased abundance of *GZMK*^+^ T cells in HGG and other types of cancer. Taken together, these data suggest that *GZMK*^+^ T cells may represent an effector subset distinct from canonically exhausted T cells and highlight this population for further study in GBM and other malignancies.

## Results

### Primary and recurrent high grade gliomas are infiltrated by a diverse repertoire of T cell states with a notable lack of exhausted phenotypes

To characterize the tumor-infiltrating T cell states and clonotypic diversity in GBM to better understand why T cells in these tumors may not be poised for licensing by checkpoint blockade immunotherapies, we performed cellular indexing of transcriptomes and epitopes by sequencing (CITE-seq) and V(D)J sequencing on an initial cohort of high grade glioma (HGG) patient samples consisting of 9 IDH-WT GBM and 1 IDH*-*mutant G4A (7 primary GBM, 2 recurrent GBM, 1 recurrent G4A) along with 5 matched peripheral blood mononuclear cell (PBMC) samples (3 matched to primary tumor, 2 matched to recurrent tumor) (Supplementary Table S1). To recognize the 2021 WHO classification of CNS tumor accurately (27), we refer to “Glioblastoma, IDH-wildtype” as “GBM” and “Astrocytoma, IDH-mutant, grade 4” as “G4A” when referencing a specific diagnosis separate from each other but use the term high grade gliomas (HGG) when discussing both diagnoses as a single cohort. The use of CITE-seq and V(D)J sequencing allowed us to intersect transcriptomic, clonotypic, and proteomic information on a single cell basis. Specifically, T cells were enriched based on CD3 expression using fluorescence activated cell sorting, stained with TotalSeq antibodies, and subsequently characterized by CITE-seq (Fig. 1A). Unsupervised clustering and uniform manifold approximation and projection (UMAP) analysis was performed on 32,320 T cells, and T cell states were identified based on the expression of both gene and protein expression (Figs. 1B and 1C, Supplementary Table S2). Detected T cell states included: CD4^+^ naїve (T_N_), central memory (T_CM_), effector memory (T_EM_), activated, *GZMK^+^* effector (T_Eff_), effector memory re-expressing CD45RA T cells (T_EMRA_), and regulatory T cells (T_reg_); CD8^+^ naїve (T_N_), *NR4A2*^lo/hi^ *GZMK^+^* effector (T_Eff_), cytotoxic resident memory (T_RM_), effector memory re-expressing CD45RA (T_EMRA_), and MAIT-like T cells; and proliferating and stress signature T cells (Figs. 1C and 1D, Supplementary Table S2, Supplementary Fig. S1, Supplementary Data S1: Sheet 1). In some cases, proteins were both highly expressed and correlated with RNA expression (e.g. CD8, Supplementary Fig. S2). In other cases, proteins were highly expressed but did not correlate with RNA expression (e.g. CD4 and CD45RA). Finally, several proteins were only moderately expressed and did not correlate well with RNA (e.g. CD27). These data highlight the utility of CITE-seq, especially with protein markers such as CD4 and CD45RA, for purposes of cell type identification. TIL derived from HGG mainly consisted of effector T cells, such as CD4^+^ T_EM_, CD4^+^ and CD8^+^ *GZMK^+^* T cells, cytotoxic T_RM_, and T_EMRA_, while T cells derived from matched PBMC samples mainly belonged to either a T_N_ or T_EMRA_ cell state (Figs. 1E and 1F).

**Figure 1:**
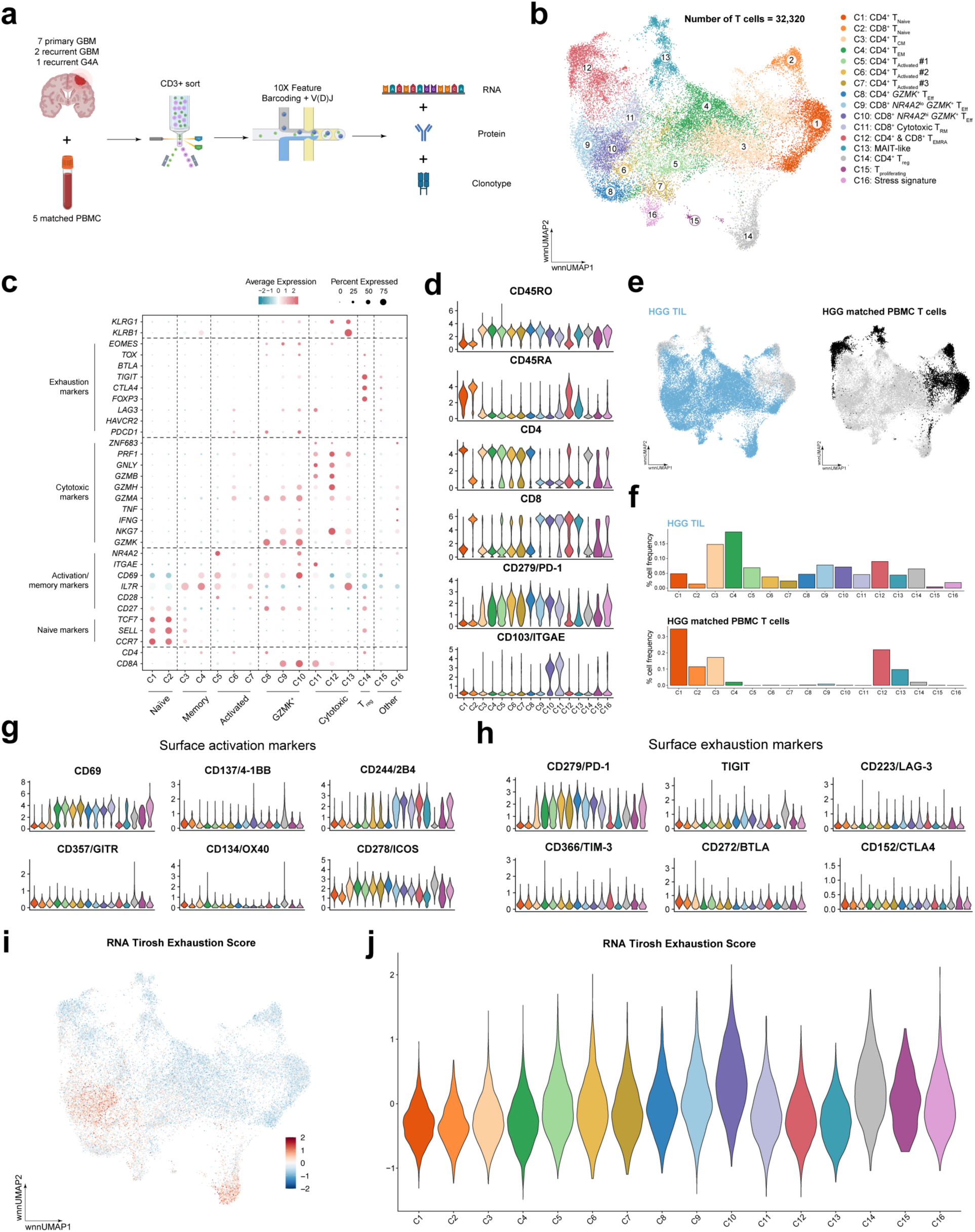
Single cell preparation and sequencing shows a diverse TIL landscape in HGG. **(A)** Illustration of single cell preparation of primary and recurrent HGG samples, consisting of both GBM and G4A, and subsequent isolation and analysis of CD3 T cells. Created with BioRender.com **(B)** UMAP visualization of 32,320 T cells in the first cohort of HGG and PBMC samples with respective cell states labeled. **(C)** Dot plot of RNA expression of select gene markers. **(D)** Violin plots of protein expression of select protein markers. **(E)** UMAP visualizations of T cells highlighted by sample type. **(F)** Bar plots of distribution of T cell states separated by sample type. **(G, H)** Violin plots of expression of select protein markers of activation and exhaustion. **(I, J)** UMAP visualization and violin plot of RNA Tirosh exhaustion score expression with each T cell state as colored in Fig. 1B.

To assess the phenotype of HGG TIL, we analyzed several cell surface protein co-stimulatory and co-inhibitory markers. High levels of CD69 and CD278/ICOS were expressed by several activated and cytotoxic T cell clusters (Fig. 1G). However, several common co-stimulatory surface receptors, including CD137/4-1BB, CD357/GITR, and CD134/OX40, were not well expressed. Finally, CD244/2B4, an NK cell receptor also expressed by T cells, was highly expressed on several CD8^+^ T cell clusters, including the *GZMK*^+^, cytotoxic T_RM_, T_EMRA_, and MAIT clusters. While PD-1 was expressed by several clusters, there were notably low levels of other exhaustion-associated markers, such as TIGIT, LAG-3, BTLA, and CTLA-4 (Fig. 1H). To determine if these PD-1 expressing clusters were represented by an exhausted transcriptional state, we applied a 28 gene expression-based exhaustion signature derived from melanoma TIL, as reported by Tirosh *et al.* (28) to our HGG TIL cohort. We scored each cell for expression of this signature (Methods) and observed a marginally higher expression in the CD8^+^ *NR4A2*^hi^ *GZMK^+^* T_Eff_ (C10) and T_regs_ (C14) clusters (Fig. 1I and 1J), while the CD8^+^ *NR4A2*^lo^ *GZMK^+^* T_Eff_ (C9) exhibited lower expression of this signature. These scores were similar between PBMC and TIL samples for each T cell cluster (Supplementary Fig. S3A). Similar findings have been reported in other GBM single cell data sets generated using different technologies, such as SmartSeq and 10X 3’ sequencing, in which GBM-infiltrating CD8^+^ T cells exhibited a weak exhaustion-associated co-inhibitory receptor expression signature (consisting of *PDCD1, CTLA4, HAVCR2, LAG3, and TIGIT*) (17,25). When using a comparable inhibitory signature as these other studies (*PDCD1, CTLA4, HAVCR2, LAG3, TIGIT,* and, *BTLA*) (17), we found relatively high expression of the inhibitory signature in several clusters, most notably the T_regs_ (C14) cluster (Supplementary Figs. S3B-S3D). Overall, the transcriptional and paired protein expression of these inhibitory receptors suggest a lack of a canonical exhausted T cell population in HGG TIL.

### Most CD8^+^ T cells within GBM TIL are GZMK^+^ and may develop separately from the T_EMRA_ cell state

Having observed the lack of a significant transcriptional exhaustion signature, we further characterized the transcriptional profile of HGG CD8^+^ T cells. A total of 10,749 CD8^+^ T cells were subsequently subsetted, reanalyzed, and visualized via UMAP (Fig. 2A, Supplementary File S1: Sheet 2). Strikingly, HGG CD8^+^ TIL were comprised more of *GZMK*^+^ T cells, either *NR4A2*^lo^ or *NR4A2*^hi^ (C9 and C10, respectively), than cytotoxic CD8^+^ T_RM_ (C11) and T_EMRA_ (C12). In PBMC, *GZMK*^+^ T_Eff_ (C9) were also present but T_N_, T_EMRA_, and MAIT-like cell states were the most prevalent cell types (Figs. 2B and 2C). Similar to *GZMK*^+^ T cells observed in aging mice, as published by Mogilenko *et al.* (29), CD8^+^ *GZMK*^+^ T cells expressed high protein levels of CD45RO, CD27, CD28, and PD-1 and low protein levels of IL7R (Fig. 2D). Transcriptionally, these cells expressed moderate levels of *TOX* and *EOMES* and low levels of *TCF7*, also previously observed by Mogilenko *et al*. in *GZMK*^+^ T cell populations derived from aged murine models (Supplementary Fig. S4). Notably, these T cells expressed high levels of *GZMK* and *GZMH* but did not express high levels of classic cytotoxic markers such as *GZMB, IFNG*, and *PRF1*. To validate whether *GZMK* is upregulated in TIL compared to PBMC on the protein level, we performed flow cytometry on prospectively collected GBM patient samples consisting of both TIL and PBMC (Figs. 2E and 2F). Gating on MAIT TCR^-^ CD45RO^+^ CD8^+^ T cells (Supplementary Fig. S5), as MAIT T cells express both GZMK and GZMB, we found that GZMK^+^GZMB^-^ and GZMK^+^GZMB^+^ T cell populations were enriched in TIL compared to PBMC (20.900 +/- 4.763% vs. 7.212 +/- 5.749%, 49.167 +/- 11.133% vs 18.688 +/- 12.760%, respectively, p<0.05, Mann-Whitney U-test). Meanwhile, GZMK^-^GZMB^+^ T cells were enriched in PBMC compared to TIL (59.500 +/- 21.318% vs. 19.167 +/- 7.920%, p<0.05, Mann-Whitney U-test). We did not observe *GZMK^+^GZMB^+^MAIT^-^*T cells in our single cell data, highlighting the limitations of single cell RNA sequencing, which does not necessarily represent the true protein-level phenotype of analyzed cells. A similar observation has been made in patients with rheumatoid arthritis, in which two major CD8^+^ T cell subsets were observed at the transcript level, one with high *GZMK^+^* expression and the other high *GZMB*^+^ expression, while three populations were observed at the protein level (GZMK^+^GZMB^-^, GZMK^+^GZMB^+^, and GZMK^-^GZMB^+^) (30).

**Figure 2:**
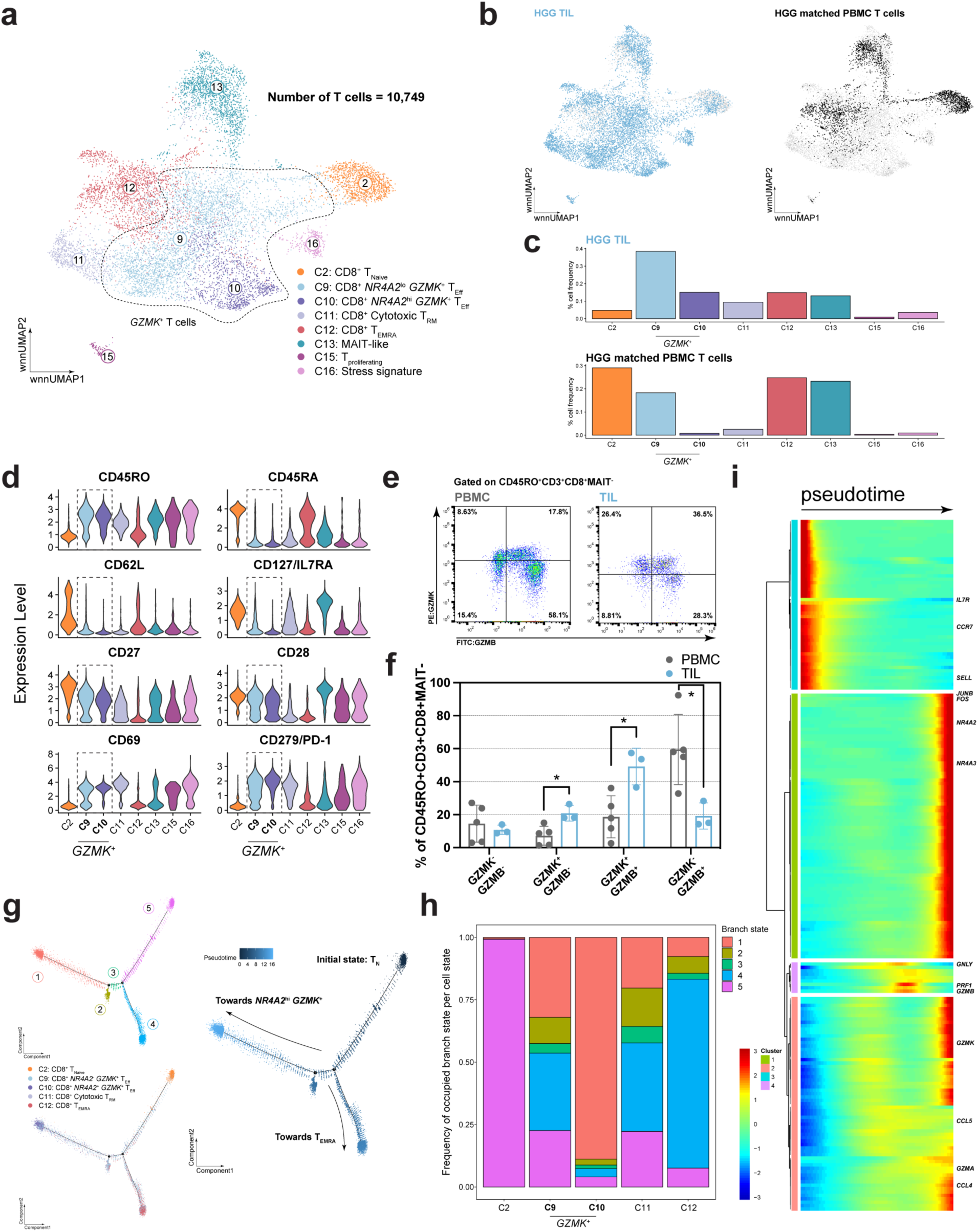
*GZMK* effector T cells represent a significant proportion of CD8 TIL and a separate development branch from T_EMRA_. **(A)** UMAP visualization of 10,749 CD8 T cells in the first cohort of HGG tumor and PBMC samples with respective cell states labeled. **(B)** UMAP visualizations of T cells highlighted by sample type. **(C)** Bar plots of distribution of T cell states separated by sample type. **(D)** Violin plots of expression of protein markers selected from Mogilenko *et al.* (29). **(E)** Representative flow cytometry plots and **(F)** cumulative bar plot of frequency of GZMK and GZMB expression gated on CD45RO^+^CD3^+^CD8^+^MAIT*^-^*cells from GBM tumor (n=3) and PBMC (n=5) samples. **(G)** Trajectory inference analysis of CD8 T cells colored by T cell state, branch state, and pseudotime. **(H)** Stacked bar plot of relative frequency of occupied branch state per cell type. **(I)** Heatmap of gene expression dynamics along pseudotime progression. Bars in **(F)** represent mean +/- SD. Significance calculated using Mann-Whitney U-test. *p<0.05.

To gain a deeper understanding of GZMK^+^ T cell development in relation to other effector subsets, we performed pseudotemporal trajectory analysis using monocle2 on select CD8^+^ T cell clusters (Fig. 2G-2I) (31). This analysis revealed several cell states and branch points beginning with T_N_ (branch state 5), then developing into *NR4A2*^lo^ *GZMK^+^* T_Eff_ and cytotoxic T_RM_, both of which were evenly distributed across several branches. From these T cell states, CD8^+^ T cells split into either T_EMRA_ (branch state 4) or *NR4A2*^hi^ *GZMK*^+^ T cells (branch state 1) (Figs. 2G and 2H). These results suggest that *NR4A2*^lo^ *GZMK^+^*and cytotoxic T_RM_ may represent a population from which terminally differentiated T cells may develop. After hierarchically clustering the top 200 differentially expressed genes plotted along the pseudotime axis, we identified several distinct gene signatures. First, genes related to the naїve T cell state, including *CCR7 and SELL*, were highly expressed at the beginning of the trajectory. In contrast, classic cytotoxic genes, such as *GNLY, PRF1,* and *GZMB*, were highly expressed towards the middle of the trajectory, while *GZMK* related genes, such as *CCL4* and *CCL5*, were highly expressed together at the end. Genes highly expressed in the *NR4A2*^hi^ *GZMK*^+^ T_eff_ subset, such as *JUNB* and *FOS* which are associated with TCR activation, clustered separately and were highly expressed towards the end of the trajectory (Fig. 2I, Supplementary File S2: Sheet 3). These results, in conjunction with the GZMK protein validation data, suggest that *NR4A2*^lo^ *GZMK*^+^ T_Eff_ cells may arise from classic cytotoxic T cells (which express exclusively *GZMB*), transition through the GZMK^+^GZMB^+^ cell state, and terminally differentiate into *NR4A2*^hi^ *GZMK^+^* T_Eff_.

### TIL are enriched for GZMK^+^ T cells in both primary and secondary brain cancers

To further validate our findings and explore whether *GZMK*^+^ T cells are enriched in other brain malignancies, we used scRNA-seq to characterize T cells isolated from a second cohort of malignant brain tumor patient samples consisting of 4 GBM samples (4 primary GBM), 2 G4A samples (1 primary and 1 recurrent), and 5 brain metastases of variable histologies (Supplementary Table S1). These samples were integrated with the scRNA-seq data from the initial cohort of GBM samples, and T cell states derived from the initial cohort were applied to the integrated data set (Fig. 3A, Supplementary File S1: Sheet 3). Notably, several new cell states were observed, specifically exhausted CD4^+^ T cells which expressed high levels of *PDCD1, LAG3, HAVCR2,* and *EOMES* (C12), while other cell states, such as the distinction between *NR4A2*^lo/hi^ *GZMK*^+^ CD8^+^ T cells, were only resolved after further subsetting (Figs. 3B-3C). These exhausted CD4^+^ T cells were mainly derived from BrMet027, a brain metastasis originating from breast cancer. Cells in this cluster scored highly for both the Tirosh RNA exhaustion signature and the RNA inhibitory signature (Fig. 3D and 3E, Supplementary Figs. S6A and S6B). These findings supported our previous observations that while *GZMK*^+^ T cells exhibited a low expression of exhaustion and inhibitory genes, they are not consistent with classically exhausted T cells according to conventional transcriptional profiles of exhaustion.

**Figure 3:**
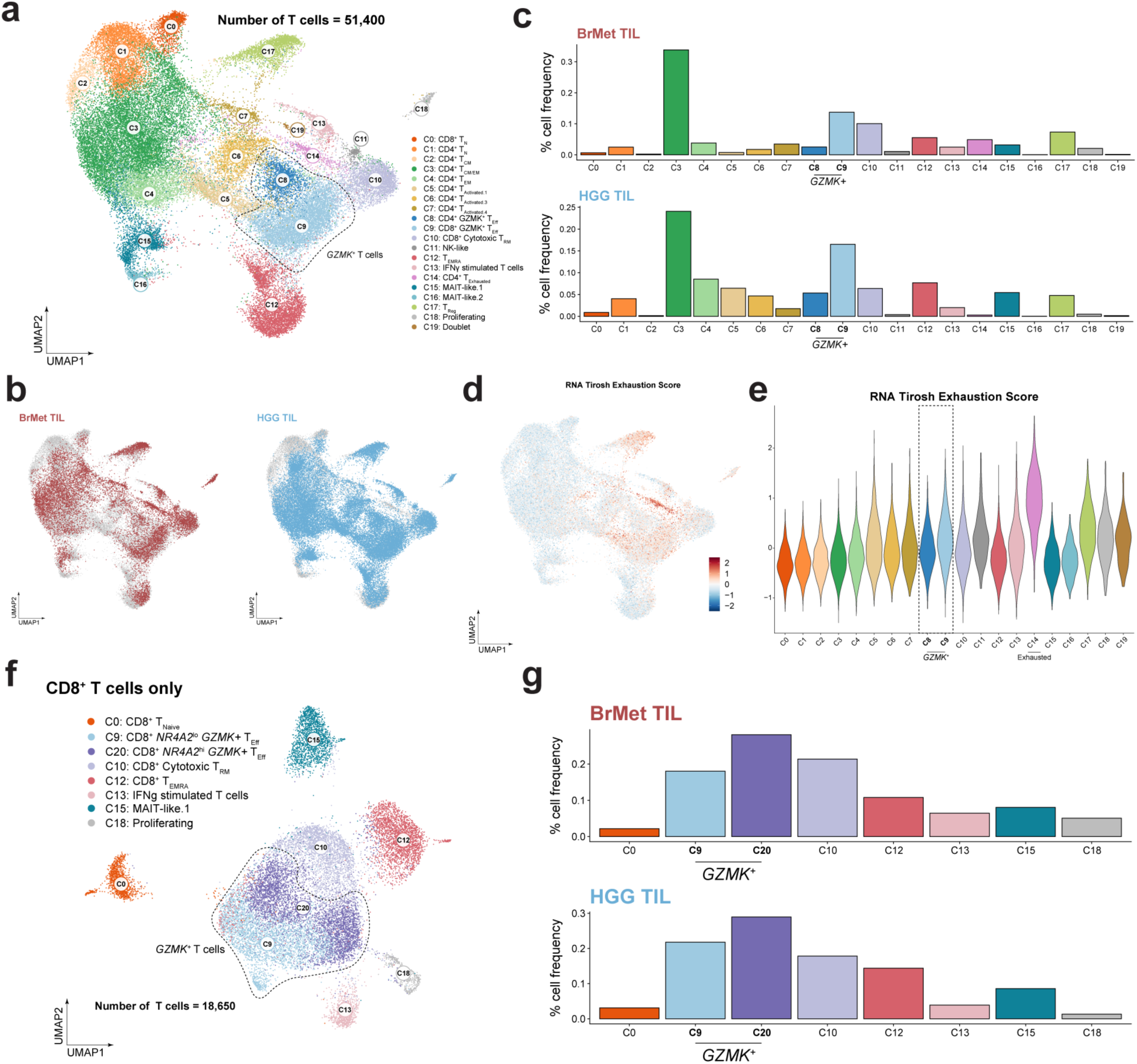
BrMet samples show exhaustion signature and enrichment of *GZMK* effector T cells. **(A)** UMAP visualization of 51,400 T cells in both the first and second cohorts of HGG, BrMet, and PBMC samples with respective cell states labeled. **(B)** UMAP visualizations of T cells with sample type highlighted. **(C)** Bar plots of distribution of T cell states separated by sample type. **(D, E)** UMAP visualization and violin plot of RNA exhaustion score expression with each cell state colored as in Fig. 3A. **(F)** UMAP visualization of 18,650 CD8^+^ T cells in both the first and second cohorts of HGG, BrMet, and PBMC samples with respective cell states labeled. **(G)** Bar plots of distribution of T cell states separated by sample type.

We observed similar proportions of *NR4A2*^lo/hi^ *GZMK*^+^ T cells with the addition of these new HGG samples compared to the initial scRNA-seq data set (16.5% vs 15.0% of total T cells, respectively), further supporting the presence of *GZMK^+^* T cells in HGG. Interestingly, BrMet samples also consisted of a similarly high proportion of *GZMK^+^*T cells (13.7%) suggesting that the presence of *GZMK^+^* T cells is not restricted to HGG but can be found in other brain malignancies (Fig. 3B and 3C). By further analyzing the CD8^+^ T cells, we resolved *NR4A2*^hi^ and *NR4A2*^lo^ *GZMK*^+^ T cells and highlighted the enrichment of *GZMK*^+^ T cells in BrMet samples with *GZMK*^+^ T cells occupying greater than 30% of the CD8^+^ T cell landscape (Figs. 3F and G, Supplementary File S1: Sheet 4). Given the overall high prevalence of *GZMK^+^*T cells in HGG and brain metastases, we next aimed to define a gene signature specific to CD8^+^ *GZMK^+^* T cells. To this end, we identified the top 25 differentially expressed genes (DEGs) by *NR4A2*^lo/hi^ *GZMK*^+^ T cells and used ToppGene (32) to perform a functional enrichment analysis (Supplementary Figure. S7, Supplementary File 3: Sheet 1). Associated biological pathways included “MHC class II protein complex assembly”, with enrichment of genes such as *HLA-DRA*, and “Lymphocyte mediated immunity”, with enrichment of genes such as *CD27* (Supplementary File 3: Sheet 2). Given *HLA-DRA* expression is associated with T cell activation, these pathways suggest that *GZMK*^+^ T cells were activated and antigen experienced. Thus, these data support a high frequency of *GZMK*^+^ T cells within GBM and BrMet CD8^+^ TIL, and that they are enriched for genes associated with T cell activation and prior antigen exposure.

### GZMK^+^ TIL are present in a broad range of cancer types

Given that *GZMK*^+^ T cells comprise a large percentage of CD8^+^ TIL, not just in GBM but also in heterogeneous BrMets, we explored whether *GZMK^+^* T cells were present in cancers outside of the central nervous system. To this end, we integrated CD8^+^ T cells from our combined HGG and BrMet data set with additional data sets comprised of CD8^+^ T cells from 1) a pan-cancer TIL data set (33), 2) a melanoma data set (34), 3) a GBM data set (17), and 4) a pancreatic ductal adenocarcinoma (PDAC) data set (35) (Fig. 4A, Supplementary Fig. S8A, Supplementary Data S1: Sheet 5) for a total of 20 different cancer types. These T cell states were annotated according to our gene signature and supported by original annotations for each respective study (Supplementary Fig. S8B). Notably, we found that *GZMK*^+^ TIL, as defined by our gene signature and supported by original annotations, were present in several tumor types and similarly separated into two distinct clusters, *NR4A2*^hi^ *GZMK*^+^ and *NR4A2*^lo^ *GZMK^+^* (Figs. 4A and 4B). The *NR4A2*^hi^ *GZMK^+^* T_Eff_ highly expressed several genes associated with TCR activation, such as *CD74, CRTAM,* and *JUNB*. Meanwhile, *NR4A2*^lo^ *GZMK^+^* T cells did not express these particular genes, suggesting these T cells were not recently activated by antigen stimulation (Supplementary Data S1: Sheet 5). A similar separation of *GZMK*^+^ T cells into these subsets has been observed in non-small cell lung cancer (NSCLC) patient samples, supporting our observations that *NR4A2* expression may delineate different activation profiles (36). Compared to our data set, the independent GBM data set (17) had higher frequencies of *NR4A2*^lo^ *GZMK*^+^ TIL and similar levels of *NR4A2*^hi^ *GZMK^+^* TIL, suggesting that our findings were generalizable across cohorts, processing protocols, and analysis pipelines. By comparing the frequency of *GZMK^+^*T cells across cancer types, we found that HGG tumors harbored some of the highest frequencies of *GZMK^+^* T cells, regardless of *NR4A2* expression. As an exhausted T cell population was identified, using both canonical gene markers and original annotations (Supplementary Fig. S8B), and clustered separately from *GZMK*^+^ T cells, we were interested in comparing the transcriptional levels of exhaustion between exhausted T cells and *GZMK*^+^ T cells. Consistent with our previous observations, *GZMK^+^* T cells (either *NR4A2*^hi^ *or NR4A2*^lo^) (C3 and C4) expressed lower levels of the RNA inhibitory and Tirosh exhaustion signatures than did exhausted T cells (C8) (p<2.2e^-16^, Mann-Whitney U-test) (Supplementary Figs. S8C and S8D). In addition, the GBM/HGG data sets had some of the lowest frequencies of exhausted T cells of the 20 cancer types analyzed (7.67% and 7.93% in our data set and the published GBM data set, respectively) (Fig. 4C). Thirteen cancer types (red bars in Fig. 4C), including our BrMet samples, had significantly higher frequencies of exhausted TIL compared to our data set, with melanoma TIL having the highest frequency (27.6%, p<0.05, Fisher’s exact test). Only two cancer types (blue bars in Fig. 4C), colorectal cancer (CRC) and pancreatic ductal adenocarcinoma (PDAC), had significantly lower frequencies of exhausted TIL than did our data set. Within the exhausted T cell population, GBM/HGG derived TIL had the lowest expression of both the RNA inhibitory and Tirosh exhaustion signatures (Supplementary Fig. S8E). These levels were statistically significantly lower when compared to all other tumor types (p<2.2e^-16^, Mann-Whitney U-test) (Supplementary Fig. S8F). This observation supported and extended, by including additional cancer types, the recent conclusion made by Naulaerts *et al.* that when compared to tumor types with high levels of canonical exhaustion, such as melanoma, GBM has few T cells that express an exhaustion signature (26).

**Figure 4:**
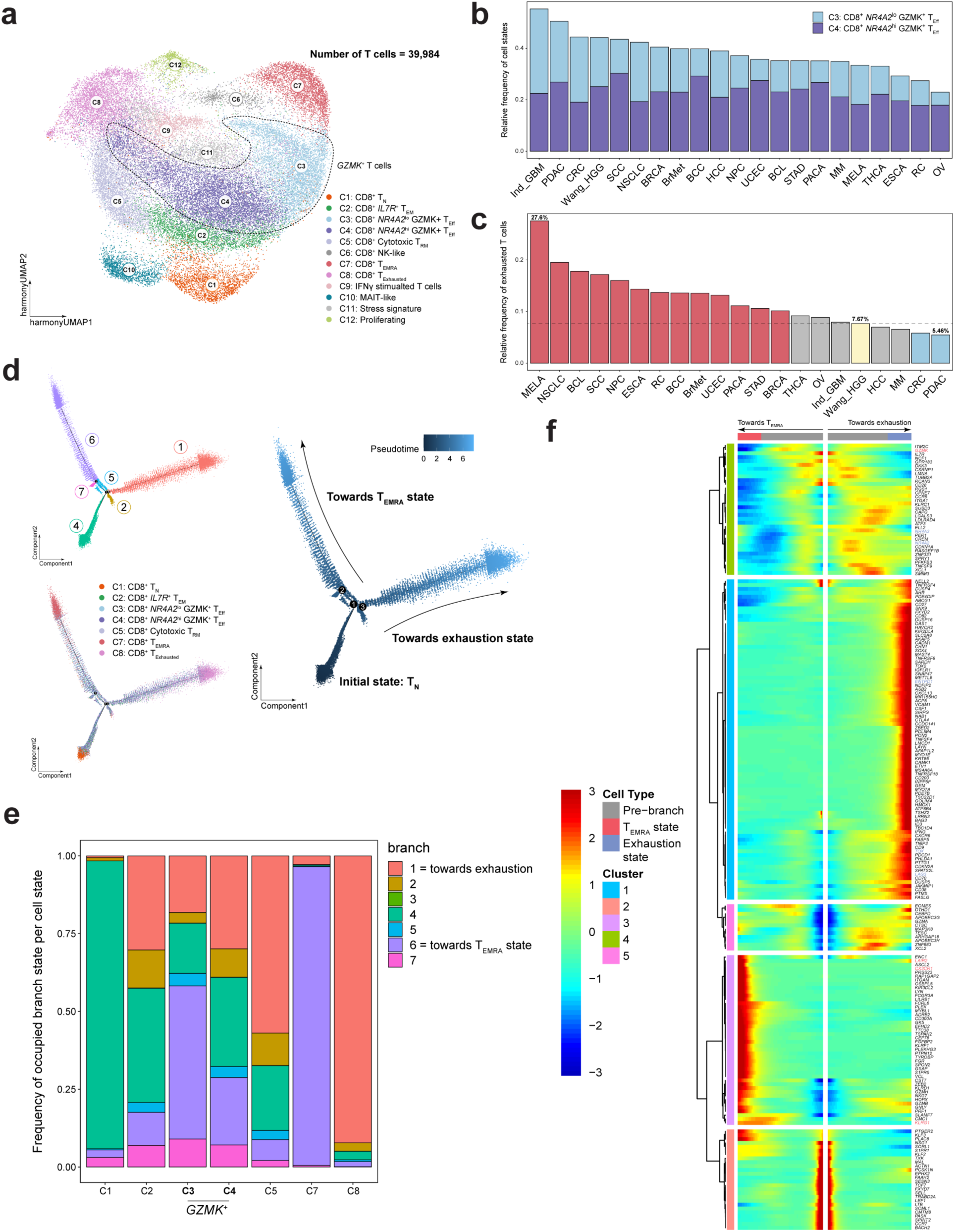
*GZMK* T cells are enriched in the many cancer types and cluster separately from exhausted T cells. **(A)** UMAP visualization of CD8 T cells derived from pan-cancer integration with respective T cell states labeled. **(B)** Stacked bar plot of relative frequency of *NR4A2*^hi^ *GZMK^+^* and *NR4A2*^lo^ *GZMK^+^* T cells among cancer types. **(C)** Bar plot of relative frequency of exhausted T cells among cancer types. The dotted lines represent the frequency of the respective T cell state in our data set (Wang *et al.*). **(D)** Trajectory inference analysis of CD8 T cells colored by T cell state, branch state, and pseudotime. **(E)** Stacked bar plot of relative frequency of occupied branch state per cell type. **(F)** Heatmap of genes differentially expressed at branch point 1. Genes upregulated on the right are genes upregulated in cells that follow the pseudotime progression to exhausted T cells while those on the left are upregulated in cells that follow the pseudotime progression to T_EMRA_. In **(C),** red bars represent cancer types with exhausted T cell frequencies that are statistically significantly increased compared to our data set. Blue bars represent cancer types with exhausted T cells that are statistically significantly decreased compared to our data set. Significance calculated using Fisher’s exact test. Red or blue bars indicate p<0.05. Ind_GBM = Independent GBM data set. In **(F)**, genes of interest are colored according to associated branch state (red represents association with the T_EMRA_ state while blue represents association with the exhausted state).

Finally, to understand how *GZMK*^+^ T cells may develop with respect to exhausted T cells, we performed pseudotemporal trajectory analysis on the integrated tumor data set. From this analysis, we observed two major terminal states branching from naїve T cells: the T_EMRA_ cell state (branch state 1) and the exhausted cell state (branch state 6) (Figs. 4D and 4E). While *NR4A2*^lo^ *GZMK^+^* T cells (C3) were more associated with the terminally differentiated cell state, *NR4A2*^hi^ *GZMK^+^* T cells (C4) were evenly split among the T_N_, T_EMRA_, and exhausted states. Consistent with previous observations, *NR4A2*^hi^ *GZMK^+^* T cells may represent a branch point from which exhausted T cells arise (36). These results differ slightly from our previous pseudotemporal trajectory analysis on isolated HGG CD8^+^ T cells suggesting *NR4A2^+^ GZMK*^+^ T cells as a terminal state, likely due to the current presence of an additional terminal differentiated state (i.e. canonical exhaustion). In addition, as expected, cytotoxic T_RM_ (C5) were mostly associated with the terminal exhausted T cell state. Next, we investigated the genes that were differentially expressed at the initial branch point which split T_EMRA_ and exhausted T cells (Fig. 4F). As expected, several genes related to exhaustion were only upregulated on the terminal end of the right branch (“towards exhaustion” in Fig. 4F), such as *TOX, LAG3, and ENTPD1*. Meanwhile, genes such as *LAIR2, KLRG1,* and *CX3CR1* were upregulated on the terminal end of the left branch (“towards T_EMRA_” in Fig. 4F), which was associated with the T_EMRA_ cell state. *GZMK* was more highly expressed at the intermediate trajectory on the T_EMRA_ branch, while genes such as *NR4A2* and *NR4A3* were more highly expressed at the intermediate trajectory on the exhausted branch. The results from these multiple pseudotemporal trajectory analyses suggest *NR4A2*^lo^ *GZMK*^+^ T cells may represent an intermediary cell state that can develop into a T_EMRA_ population in the absence of antigen stimulation but alternatively, can differentiate into an exhausted T cell population via a *NR4A2*^hi^ *GZMK*^+^ T cell state with persistent antigen stimulation.

### GZMK^+^ T cells within the tumor microenvironment are clonally expanded

Several prior studies have demonstrated that subsets of T cells within GBM are clonally expanded (10,17). We therefore investigated the relationship between *GZMK* expression and both clonotype diversity and expansion using single cell V(D)J sequencing. Of the 18,650 CD8*^+^* T cells sequenced, 11,360 T cells (60.9%) contained paired αβ TCRs (Fig. 5A). T cell expansion states were defined as follows: TCRs with a count of 1 were labeled as singlets, with a count of 2 as doublets, with a count greater than 2 but less than or equal to 50 as expanded, and with a count greater than 50 as hyperexpanded. Among all T cell states, as expected, T_N_ (C0) were mostly single clones while around 50% of *NR4A2*^lo^ *GZMK*^+^ (C9), *NR4A2*^hi^ *GZMK^+^*(C20), and cytotoxic T_RM_ (C10) clusters and up to 75% of the T_EMRA_ cluster (C12) were either expanded or hyperexpanded (Fig. 5B). When investigating the associated T cell states of the top 50 clonally expanded TCRs for each sample type, 2 of the top 3 expanded TCRs in both PBMC and HGG samples were invariant MAIT TCRs. Of the total number of cells associated with these clonally expanded TCRs, 34.1% and 11.9% were associated with the MAIT T cell state in PBMC and HGG, respectively. In PBMC, the T_EMRA_ state was the most frequent non-MAIT T cell state associated with these expanded TCRs (35.1%). *NR4A2*^lo^ *GZMK^+^, NR4A2*^hi^ *GZMK^+^,* and cytotoxic T_RM_ only occupied 9.80%, 11.6%, and 8.97% of the total expanded T cell landscape, respectively. In contrast, within HGG TIL, *GZMK*^+^ T cells comprised the majority of clonally expanded T cells with a relatively equal proportion of both *NR4A2*^lo^ and *NR4A2*^hi^ GZMK T cells (24.4% and 29.1%, respectively), and were significantly enriched compared to respective populations in PBMC (9.80% and 11.6%, respectively) (p<0.05, Fisher’s exact test). Meanwhile, cytotoxic T_RM_ and T_EMRA_ occupied only 9.86% and 20.4% of the clonally expanded T cell landscape in HGG samples. Finally, in BrMet samples, cytotoxic T_RM_ occupied the greatest proportion of clonally expanded T cells (29.2%) followed by *NR4A2*^lo^ and *NR4A2*^hi^ GZMK T cells (18.2% and 28.4%, respectively) (Fig. 5C). These results suggested that more *GZMK^+^* T cells are associated with the most expanded TCRs compared with any other T cell subsets within the brain tumor microenvironment. Moreover, these enriched clonotypes were shared by both *NR4A2*^hi^ and *NR4A2*^lo^ *GZMK*^+^ T cells, supporting our hypothesis that *NR4A2*^hi^ *GZMK*^+^ T cells may develop from *NR4A2*^lo^ *GZMK*^+^ T cells. Interestingly, between tumor types, *GZMK*^+^ T cells occupied a greater proportion of the total T cell landscape in HGG compared to BrMet, with cytotoxic T_RM_ more abundant in the latter.

**Figure 5:**
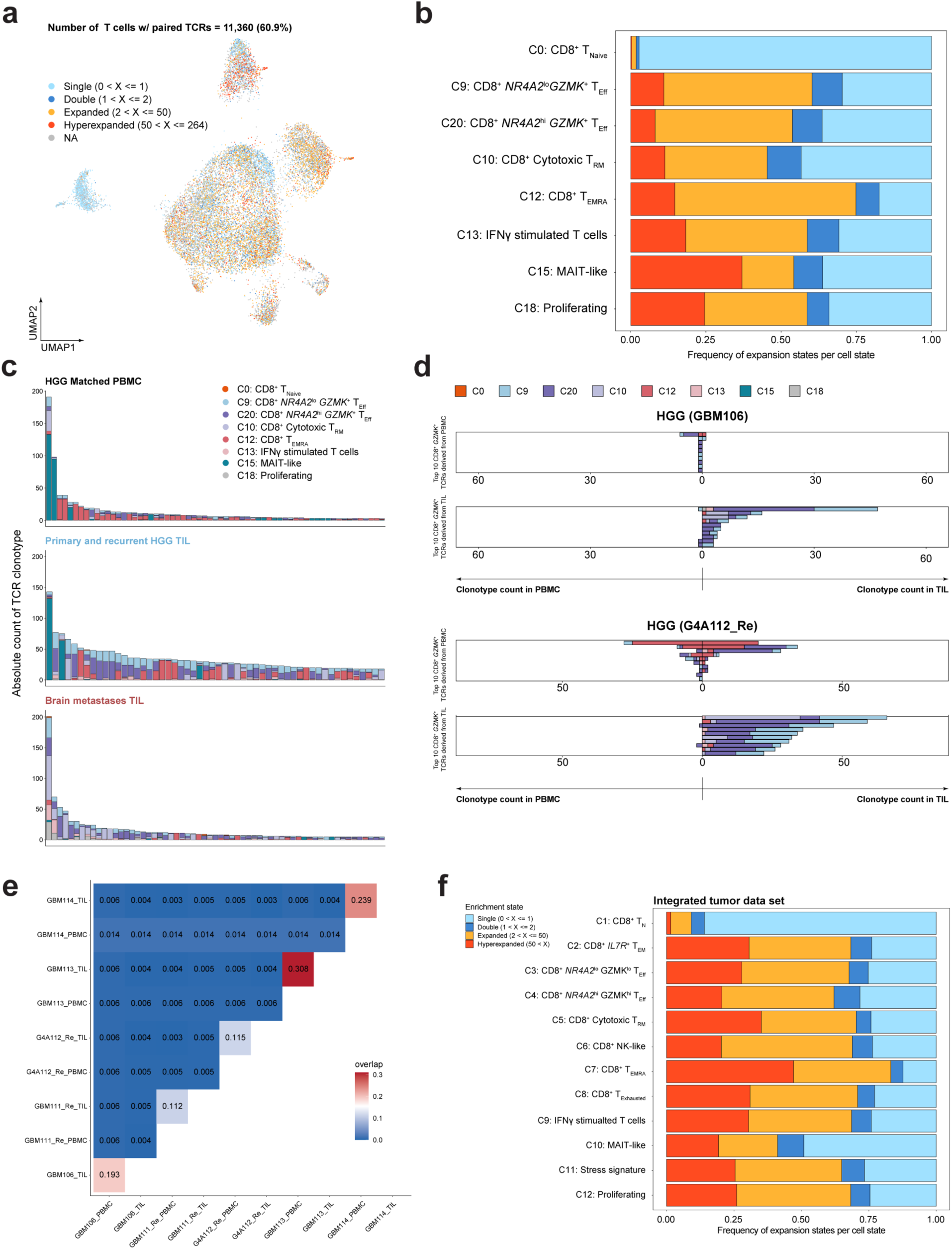
Expanded *GZMK* T cells are enriched in HGG TIL and are specific to the tumor site, and not matched peripheral blood. **(A)** UMAP visualization of CD8 T cells from Fig. 3E with respective TCR clonotype enrichment states labeled. **(B)** Stacked bar plot of distribution of TCR clonotype enrichment states by T cell state. **(C)** Distributions of the top 50 expanded TCR clonotypes colored by respective T cell states from Fig. 3E and separated by sample type. **(D)** Representative double sided bar plots representing the absolute count of the top 10 CD8^+^ *NR4A2*^lo/hi^ *GZMK^+^* T_Eff_ (C9 & C20) TCRs from matched PBMC (top) or TIL (bottom) samples within the total population of CD8 T cells in paired samples. T cell states associated with each TCR are colored as in Fig. 3E. **(E)** Heatmap of overlap coefficient for quantification of CD8^+^ T cell clonotype overlap among all five paired PBMC and TIL samples. **(F)** Stacked bar plot of distribution of TCR clonotype enrichment states by T cell state in the integrated pan-cancer data set.

To determine if expanded TCRs expressed by *GZMK*^+^ T_Eff_ were shared by other T cell states, we investigated the clonotype count, distribution, and associated T cell states of the top 10 expanded non-MAIT TCRs associated with *GZMK^+^* T_Eff_. Pairwise analysis was performed on autologous, matched PBMC and TIL from two patients (Fig. 5D). In the case of GBM106, the top 10 expanded TCRs associated with the *GZMK^+^* T cell state in GBM106 PBMC were minimally expanded, with only one TCR detected more than once. This TCR was detected in a total of six PBMC associated T cells: five *GZMK*^+^ T cells and one T_EMRA_ cell. Overall, these same 10 TCRs were either not present in the TIL or detected with a single count. In comparison, the top 10 expanded TCRs associated with the *GZMK^+^* T cell state in GBM106 TIL were detected at a much greater frequency. For example, the most highly expanded TIL clone was detected in 47 cells, with 17 and 27 of these cells associated with the *NR4A2*^lo^ or *NR4A2*^hi^ *GZMK^+^* T cell state, respectively. Notably, all TCRs were mostly, if not entirely, associated with either the *NR4A2*^lo/hi^ *GZMK^+^* T cell state or shared with the cytotoxic T_RM_ state. Importantly, these top expanded TCRs in the TIL were either undetected or detected only once in the matched PBMC sample. In the case of patient G4A112_Re PBMC, the top 10 expanded TCRs associated with the *GZMK^+^* T cell state were detected at higher frequencies than TCRs in GBM106 PBMC. However, these expanded TCRs were mostly associated with the T_EMRA_ state, as seen with the top two most expanded TCRs. These TCRs were detected in TIL and either exclusively expressed by the T_EMRA_ state or shared with the *NR4A2*^hi/lo^ *GZMK^+^*T cell state, suggesting that T_EMRA_ and *GZMK^+^* T cell states may also share a developmental trajectory. Given the presence of these expanded TCRs in both PBMC and TIL, they may reflect infiltration of non-tumor-specific bystander T cells. Meanwhile, the top 10 expanded TCRs associated with the *GZMK^+^* T cell state in G4A112_Re TIL were heavily expanded solely in the TIL and not the matched PBMC sample. Similarly, these TCRs were largely specific to the *NR4A2*^lo/hi^ *GZMK*^+^ T cell state, although several TCRs, the most expanded in particular, were also shared by cytotoxic T_RM_. However, in TIL, we did not detect shared expression of specific TCRs between T_EMRA_ and *GZMK^+^* T cells. The observation that *GZMK*^+^ TCRs were selectively expanded in TIL compared to PBMC was representative of the general clonotypic landscape of all matched samples as paired TIL and PBMC samples had low overlap (Fig. 5E). Given that T cells with the same TCRs are derived from the same progenitor cell, these results overall support our pseudotemporal trajectory analysis suggesting that cytotoxic T_RM_ and *NR4A2*^lo/hi^ *GZMK*^+^ T cells lie along the same developmental trajectory in the tumor microenvironment.

To determine whether expansion of *GZMK*^+^ T cells was specific to brain tumors, we investigated the TCR expansion states in our integrated tumor data set. Of the 20 tumor types examined, 13 contained V(D)J data that allowed for quantification of expansion states (Supplementary Figs. S8G and S8H). Notably, we observed that within the integrated tumor data set, both *NR4A2*^lo^ and *NR4A2*^hi^ *GZMK*^+^ T cells (C3 & C4) were expanded, with greater than 60% of cells either expanded or hyperexpanded (Fig. 5F).

From these results, we concluded that expanded *GZMK*^+^ TCRs were exclusively expanded in the tumor setting compared to matched blood; within the tumor microenvironment, they were mostly expressed by either *GZMK*^+^ T cells or cytotoxic T_RM_. Furthermore, expansion of *GZMK*^+^ T cells was not specific to brain tumors but occurred in many different cancer types arising from diverse anatomical sites.

### CD8^+^ GZMK^+^ TCRs are not detectable following ex vivo expansion

Adoptive cellular transfer of *ex vivo* expanded TIL derived from patient tumor specimens has been shown to be a promising immunotherapeutic approach in several cancer types (37,38). As antigen-specific T cells are enriched within the tumor microenvironment, isolated tumor specimens represent an ideal source of tumor-specific effector T cells for such immunotherapies. However, as many terminally differentiated T cell states lose capacity for self-renewal, it is unclear which effector T cell states are truly expanded from these *ex vivo* techniques. Recently, TCRs associated with *GZMK^+^* TIL derived from PDAC patient samples have been shown to sustain similar frequencies following *ex vivo* culture, suggesting *GZMK^+^* TIL retain proliferative potential (35). As a result, we were interested in assessing if these putative antigen-experienced, clonally expanded CD8^+^ *NR4A2*^lo/hi^ *GZMK*^+^ T_Eff_ derived from HGG patient samples were similarly capable of *ex vivo* expansion for potential therapeutic considerations. To this end, TIL from 6 different HGG patient samples were expanded in a gas permeable rapid expansion (G-Rex) culture system and subsequently analyzed via bulk TCR CDR3 beta-chain sequencing using the immunoSEQ platform (Fig. 6A). Among all paired samples, we observed an increase in overall clonal expansion following *ex vivo* culture represented by an increase in the occupied repertoire space by the topmost expanded TCRβ chains (Fig. 6B). However, when analyzing the TCR repertoire of each sample type, we found that cultured TIL TCR clonotypes were not representative of either D0 TIL or D0 PBMC. Such results are shown by representative alluvial plots of the top 20 TCRβ chains of D0 TIL, D25 cultured TIL, and D0 PBMC samples derived from patient sample GBM111_Re (Supplementary Fig. S9A). When investigating specifically the top 20 TCRβ chains of D0 TIL, we observed minimal enrichment of such TCRβ chains in D25 cultured TIL and some enrichment of such TCRβ chains in D0 PBMC. The inverse of such results is observed when investigating the top 20 TCRβ chains of D25 cultured TIL. These results were summarized across all samples with the use of the Morisita’s overlap index, a measurement of the overlap among different data sets (Supplementary Fig. S9B), and similar results have been previously reported (39). Specifically analyzing the expansion status of TCRs associated with CD8^+^ *GZMK^+^*T cells, we matched bulk TCRβ chains with those from CD8^+^ *GZMK*^+^ T cells within the single cell TCR data sets. In two representative samples, GBM111_Re and GBM113, we observed that these matched TCRβ chains were present in D0 TIL but were either not detected or unexpanded in cultured TIL, aside from one TCRβ chain which was expanded in cultured GBM113 TIL (Figs. 6C and 6D). Overall, these results suggested that the clonally expanded *GZMK*^+^ TIL present in the initial HGG microenvironment have limited expansion potential *ex vivo* using conventional expansion conditions. This potential loss of proliferative capacity, supported by low *TCF7* expression (Supplementary Fig. S4), may also be HGG specific as expanded *GZMK^+^* TIL derived from PDAC patient samples have been shown to continuously expand *ex vivo* (35).

**Figure 6:**
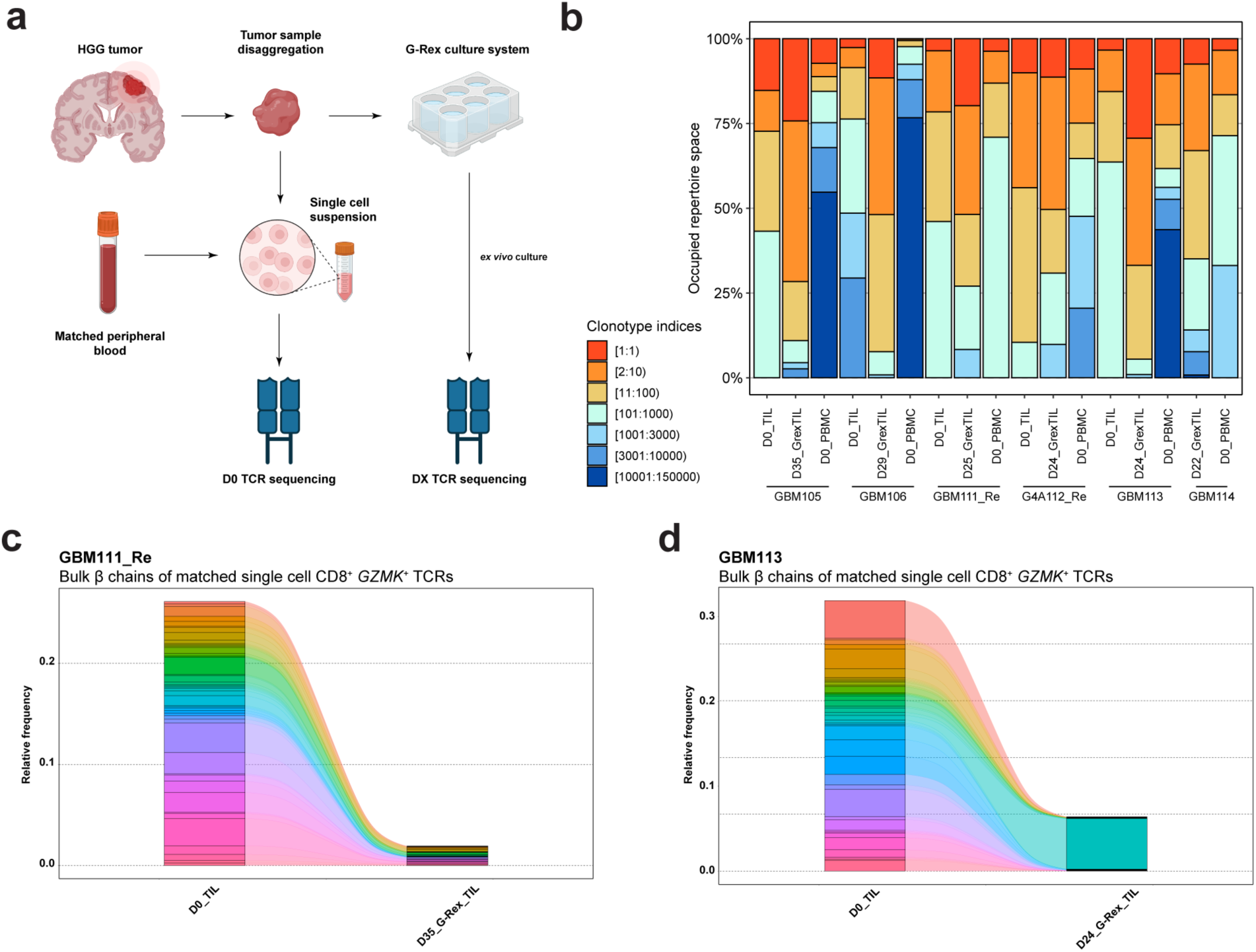
TIL expanded *ex vivo* are not representative of the initial TCR landscape. **(A)** Schematic of *ex vivo* growth of HGG TIL in G-rex and subsequent downstream analysis. Created with BioRender.com. **(B)** Bar plot of frequency of clonotype expansion states among various samples. **(C, D)** Alluvial plots highlighting the overlap between matched sample types of the β chains matched to single cell CD8^+^ *GZMK*^+^ TCRs in D0_TIL.

## Discussion

In this study, we provided a combination of CITE-seq and scRNA-seq to explore the proteogenomic landscape of T cells within primary and recurrent HGG, including GBM and G4A, matched peripheral blood (PBMC), and brain metastases (BrMet). With paired single cell V(D)J sequencing, several observations emerged. First, using protein markers, we labeled T cell states that were otherwise unidentifiable by scRNA-seq alone and characterized the cell surface protein phenotypes of T cell states. Second, we observed an unexpected lack of canonical exhaustion markers at both the RNA and protein level, which is supported by other single cell analyses of GBM TIL, but had not been thoroughly compared to many other cancer types (17,39). Third, *GZMK*^+^ T cells comprised the largest percentage of CD8^+^ TIL within malignant brain tumors and were enriched in the tumor microenvironment compared to peripheral blood. Interestingly, this subset was further defined by the expression level of *NR4A2*. Fourth, through pseudotemporal ordering analysis, we inferred that *NR4A2*^hi^ *GZMK*^+^ T cells may represent a terminal developmental state derived from naїve CD8^+^ T cells and transitioning through *NR4A2*^lo^ *GZMK^+^*T cells. Fifth, CD8^+^ *GZMK*^+^ TIL comprised the most clonally expanded T cells relative to peripheral blood as well. Finally, when we integrated our data set, including both HGG and BrMet samples, with those from 18 other cancer types, we observed that *GZMK*^+^ TIL were found across all cancer types at varying frequencies, with higher frequencies in HGG. Taken together, these data further refine our understanding of the T cell populations within malignant brain tumors and across many other cancer types.

The comparative integrated analysis with additional cancer data sets harboring T cells with high-level exhaustion signatures, such as those found in melanoma, highlighted the profound differences in T cell exhaustion states between HGG and other cancers. In the overall T cell landscape of our data set, we observed high levels of inhibitory receptor PD-1 protein and *PDCD1* RNA, moderate levels of TIGIT protein and RNA, and low levels of LAG-3, TIM-3, and BTLA protein and RNA. When applying an exhaustion gene signature based upon 28 genes upregulated in functionally exhausted T cells derived from melanoma (28), we found slightly elevated expression of this signature in *GZMK^+^* T_Eff_ and T_reg_ clusters. Compared with 19 other cancer types, including BrMets and cancers known to harbor high levels of exhausted T cells, HGG, and in particular GBM, was infiltrated by some of the lowest frequencies of exhausted T cells. Our conclusions also support and expand the findings of a recent study by Naulaerts *et al*. which demonstrated that GBM TIL did not express strong exhaustion signatures (26). The low expression of canonical exhaustion markers by T cells, coupled with the low overall frequency of T cell infiltration in GBM, could help explain the lack of efficacy of anti-PD-1/PD-L1 therapies in patients with GBM (3,4). Additional mechanisms driving T cell inhibition or dysfunction, such as activation of CD161 (17) or IL-10 release (20) by myeloid cells, could be more therapeutically compelling targets.

Our focus turned to *NR4A2*^lo/hi^ *GZMK^+^* T_Eff_ that highly expressed *GZMK*, but did not express classic cytotoxic genes, such as *GZMB, GNLY,* and *PRF1*. While this T cell transcriptional state has been previously described in several single cell studies that encompass many different cancer histologies (26,30,35,40–44), several findings herein are novel. We found that *GZMK*^+^ T cells occupied a significant proportion of T cells in HGG both within our own data set and another published study, especially in contrast to other tumor types and matched blood and are further characterized based on *NR4A2* expression. Moreover, these data support a developmental trajectory in which *GZMK^+^* T cells may arise from classic cytotoxic T cells that acquire *NR4A2* expression. Strikingly, tumor derived *GZMK^+^* T cells were clonally expanded, whereas TCR matched *GZMK*^+^ T cells derived from blood were either not expanded or not detectable in matched blood, suggesting *GZMK*^+^ T cells either homed to the tumor site or developed *in situ*. While *GZMK*^+^ T cells were similarly abundant in metastatic brain samples, cytotoxic T_RM_ were more clonally expanded compared to *GZMK*^+^ T cells. The selective enrichment of clonally expanded *GZMK*^+^ T cells in the tumor microenvironment may suggest that these T cells are antigen-experienced and potentially tumor-specific. Further functional studies are needed to explore this possibility and identify the cognate antigen targets. Interestingly, a prior study in lung cancer demonstrated that *GZMK*^+^ T cells were equally clonally expanded in the blood and in lung tumors and suggested that further differentiation occurred in the tumor microenvironment (45). This observation, when compared to our findings, raises the intriguing possibility that T cell developmental fates may vary amongst tumor types and merits additional examination.

As observed in the initial cohort of HGG and PBMC samples, when integrating our data set with several other cancer types, we noted two distinct populations of *GZMK*^+^ T cells: *NR4A2*^hi^ and *NR4A2*^lo^ *GZMK*^+^ T cells. *NR4A2*^hi^ *GZMK^+^* T cells were enriched for many genes associated with, and downstream of, TCR activation, whereas these genes were significantly downregulated in the *NR4A2*^lo^ *GZMK*^+^ T cells. *NR4A2,* which encodes NR4A2 and also known as NURR1, appears to perform several functions. In T cells, *NR4A2* has been associated with resident memory differentiation (46,47). In addition, the *NR4A* family has been associated with modulating T cell function, as triple knock out (TKO) of the *NR4A1, NR4A2,* and *NR4A3* transcription factors significantly increased the effector function of CAR T cells *in vivo* (48). Specifically, TKO of *NR4A1-3* in CAR T cells promoted tumor regression and decreased expression of several inhibitory receptors while increasing the expression of effector genes. Furthermore, recent studies of the double knock out (DKO) of the *TOX* transcription factor family in murine models revealed a cross-regulatory relationship between the *TOX* and *NR4A* transcriptional families (49). Given that the most expanded *GZMK^+^* TCRs were shared by both *NR4A2*^hi^ and *NR4A2*^lo^ transcriptional states, these results suggest that a switch in transcriptional state occurs. While we hypothesize that *NR4A2*^hi^ *GZMK^+^* T cells develop from *NR4A2*^lo^ *GZMK^+^*T cells, further study is required to understand the developmental relationships and trajectories of these T cell subsets.

As adoptive T cell therapy (ACT) has become an increasingly developed technology and expanded *GZMK^+^*TIL from PDAC patient samples have been shown to further expand *ex vivo* (35), we were interested in investigating whether clonally expanded *GZMK*^+^ TIL from HGG would likewise expand *ex vivo*. While TIL were expanded in this system, with enrichment of several clonotypes, they recapitulated neither the initial TIL nor PBMC clonotype landscape. Similar observations have been made by another group in which the TCR landscape of expanded TIL from GBM did not represent the original TIL TCR landscape (39). In our study, we additionally showed that *GZMK*^+^ TIL likewise demonstrated this pattern. Specifically, bulk TCRβ chains matched to single cell TCRs associated with the *GZMK*^+^ T cell state were expanded in initial HGG TIL but following *ex vivo* culturing were either minimally expanded or not detected. These results indicated that first, the use of current T cell expansion systems will likely need to be modified for further expansion of the most clonally expanded TIL present within the HGG tumor microenvironment. If these populations, which include *GZMK^+^* subsets, possess tumor-specific T cells, then an alternative approach to generating large numbers of these expanded clonotypes is required for ACT to be particularly effective for GBM treatment. Second, these results highlighted that *GZMK*^+^ T cells initially present in the HGG tumor microenvironment may not be capable of further proliferation. Given expanded *GZMK*^+^ TIL from PDAC have been shown to further expand *ex vivo* (35), this loss of proliferative capacity may be glioma specific and contribute to immune evasion by HGG tumor cells. Further work is needed to understand the reversibility of this observation.

The emerging identification of both RNA and protein expression of GZMK within TIL raises the question of what the function of GZMK expression may be in these biological settings. Several studies have revealed the various functions of GZMK, both intracellular and extracellular. Intracellularly, GZMK has been shown to induce both caspase-dependent and -independent cell death *in vitro*, in addition to non-cytotoxic functions, such as inhibiting viral replication (50–54). Extracellularly, GZMK has been implicated in promoting a pro-inflammatory cytokine response (54,55). Notably, a recent study by Mogilenko *et al.* investigated the effects of age on the immune system and discovered the enrichment of *GZMK*^+^ T cells in older mice (29). These *GZMK*^+^ T cells developed in response to unknown extrinsic factors and expressed several exhaustion markers. Interestingly, GZMK was shown to induce a pro-inflammatory response, specifically stimulating production of IL6 and CCL2, both components of the senescence-associated secretory phenotype (SASP), by murine fibroblasts *in vitro*. In this way, *GZMK*^+^ T cells may contribute to age-related inflammation. Meanwhile, recent work showing that *GZMK* expression is affected by chronic antigen stimulation in both murine T cells or human CAR T cells underscores a potential role for its expression as a biomarker of antigen experience and also highlights a potentially distinct lineage separate from canonical exhaustion (56,57). In murine models, *GZMK* was initially expressed at day 0 (D0), downregulated at D4, and upregulated at D7. Meanwhile, *GZMB* continuously increased following chronic antigen stimulation. Moreover, human CAR T cells harboring the 4-1BB signaling domain were dominated by a *GZMK*^+^ transcriptional profile following 15 days of antigenic stimulation. In contrast, human CAR T cells containing the CD28 signaling domain were only enriched by a canonically exhausted transcriptional profile. These results suggest that the *GZMK*^+^ transcriptional profile, perhaps influenced by distinct signaling inputs, may represent a functional state that is distinct from the canonical exhaustion state. In our own study, we performed functional enrichment analysis on the top 25 DEGs by CD8^+^ *GZMK*^+^ T cells and found several pathways suggesting these T cells were activated. In conjunction with the selective expansion of these T cells in the tumor microenvironment and lack of exhaustion markers, we hypothesize that *GZMK*^+^ T cells are tumor specific and potentially functional. Finally, further work is also needed to understand how *GZMK^+^*T cells may interact with other immune cell types within the microenvironment. Specifically, a recent study on non-metastatic colorectal tumors found that neutrophils were associated with *GZMK^+^* T cells in the tumor microenvironment, which eventually promoted tumor relapse (58).

Taken together, our findings provide a detailed analysis of the T cell states in HGG, consisting of both GBM and G4A, metastatic brain tumors, and matched PBMC samples. Given the unique and enriched clonal expansion of *GZMK^+^*TIL, investigation of the antigen specificity of this T cell subset as well as how best to modulate, such as by licensing or expanding, this population to maximally augment its effector functions in patients with brain tumors may hold clinical relevance.

## Materials and Methods

### Patient recruitment and sample collection

Adult patients undergoing neurosurgical intervention at Barnes-Jewish Hospital and Massachusetts General Hospital were screened. Selection criteria included (1) age > 18 years and (2) presence of either primary or recurrent glioblastoma or high-grade glioma with clinical indications for surgical resection. Informed consent was obtained from patients prior to surgery following the Washington University School of Medicine Institutional Review Board (IRB) Protocol #202107071, Washington University School of Medicine IRB Protocol #2011110011 and #202101040, the Dana Farber/Harvard Cancer Center IRB Protocol #10-417 and secondary Massachusetts General Hospital IRB Protocol #2022P001982, and the Massachusetts General Hospital IRB Protocol #2022P000955. Following surgical resection, specimens were placed in normal saline and maintained on ice. Characteristics of patients are summarized in Supplementary Table S1. All procedures and experiments were performed in accordance with the Helsinki Declaration.

### Clinical Sample Processing: Single Cell RNA Sequencing

Tumor samples were processed as previously reported (10). Cohort 1 samples were stained with anti-CD45, anti-CD3, anti-CD11b, Zombie NIR Viability Dye, and TotalSeq-C antibodies (Supplementary Table S3). Live CD3^+^CD11b*^-^*single cells were sorted by FACS on a BD FACSAria II (BD Biosciences) and a single cell suspension was submitted for sequencing. Cohort 2 samples were similarly stained with anti-CD45, anti-CD3, anti-CD11b, and Zombie NIR Viability Dye. Live CD45^+^ single cells were sorted by FACS on a BD FACSAria II (BD Biosciences) and a single cell suspension was submitted for sequencing.

Peripheral blood mononuclear cells (PBMCs) were separated through Ficoll-Paque PLUS density gradient (GE). The resulting buffy coat was collected, stained, and submitted for sequencing.

### Clinical Sample Processing: Flow Cytometry

Tumor samples were collected from the operating room on ice and initially weighed. Samples were separated into aliquots up to 750 mg, as recommended by the Brain Tumor Dissociation Kit (Miltenyi Biotec). Aliquots were macerated on ice and disaggregated using the Brain Tumor Dissociation Kit (Miltenyi Biotec) on the gentleMACs Octo Dissociator with Heaters (Miltenyi Biotec). Samples then underwent Percoll (GE Healthcare Life Sciences) density gradient centrifugation to remove myelin contamination. The resulting pellets then underwent RBC lysis with ACK Lysis Buffer (Lonza Biosciences) and were passed through 70 micron filters. The tumor samples were then stained with anti-CD45RO, anti-CD3, anti-CD4, anti-CD8, anti-GZMK, anti-GZMB, anti-TCR Vα7.2, and Zombie Aqua Viability Dye (Supplementary Table S3) and analyzed on a Cytoflex LX (Beckman Coulter) or Sony MA900 (Sony).

### Single Cell Library Preparation (CITE-seq/scRNA-seq/VDJ-seq)

Single cell gene expression and V(D)J libraries were prepared using the Chromium Next GEM Single Cell 5’ Reagents Kits v1 and Chromium Single Cell V(D)J Reagent Kits v1.1 (10X Genomics), respectively. cDNA was prepared after the GEM generation and barcoding, followed by the GEM-RT reaction and bead cleanup steps. Purified cDNA was amplified for 10-14 cycles before being cleaned up using SPRIselect beads. Samples were then run on a Bioanalyzer to determine the cDNA concentration. V(D)J target enrichment (TCR) was performed on the full length cDNA. Gene Expression, Enriched TCR and Feature libraries were prepared as recommended by the 10x Genomics Chromium Single Cell V(D)J Reagent Kits (v1 Chemistry) user guide with Feature Barcoding technology for Cell Surface Protein and Immune Receptor Mapping user guide, with appropriate modifications to the PCR cycles based on the calculated cDNA concentration. For sample preparation on the 10x Genomics platform, the Chromium Single Cell 5’ and Gel Bead Kit v2, 16 rxns (PN-1000006), Chromium Single Cell A Chip Kit, 48 rxns (PN-1000152), Chromium Single Cell V(D)J Human TCR Enrichment Kit (PN-1000005), Chromium Single Index Kit T, 96 rxns (PN-1000213), Chromium Single Cell 5’ Feature Barcode library Kit, 16 rxns (PN-1000080) and Single Index Kit N Set A, 96 rxns (PN-1000212) were used. The concentration of each library was accurately determined through qPCR utilizing the KAPA library Quantification Kit according to the manufacturer’s protocol (KAPA Biosystems/Roche) to produce cluster counts appropriate for the Illumina NovaSeq6000 instrument. Normalized libraries were sequenced on a NovaSeq6000 S4 Flow Cell using the XP workflow and a 151×10×10×151 sequencing recipe according to the manufacturer protocol. A median sequencing depth of 50,000 reads/cell was targeted for each Gene Expression library and 5,000 reads/cell for each V(D)J and Feature library.

### Single Cell Sequencing Analysis

Raw sequencing data generated from 5’ gene expression libraries from both cohorts 1 and 2 were processed with the CellRanger pipeline (10X Genomics, default settings, version 3.0.1) and mapped onto a human genome reference (GrCh38-2020-A). Downstream analysis was performed using the Seurat R package (59). Initial filtering excluded cells that had any of the following: a nFeature count less than 500, mitochondrial reads greater than 10%, or a nCount value greater than the 93rd percentile of each individual sample. All TCR related genes were then removed. Following this, doublets were initially identified using the DoubletCollection R package (60) in which several doublet packages were used to label doublets within each sample. The specific packages used included scds (61), scDblFinder (62), doubletCells (63), DoubletFinder (64), and Scrublet (65). Cells labeled as a doublet by three or more packages were then removed.

For samples within Cohort 1, which contained both RNA and antibody-derived tag (ADT) (i.e. or protein), single cell data, the RNA assay was first integrated using the reciprocal PCA (RPCA) method. Principal component analysis (PCA) on the integrated data set was then performed and the optimal number of principal components (PCs) was determined based upon elbow plots, jackstraw resampling, and PC expression heatmaps. Next, the ADT assay was similarly integrated using the RPCA method. The RNA and ADT assays were then integrated together using the Weighted Nearest Neighbor Analysis (WNN) (66). Dimensionality reduction and visualization were performed with the uniform manifold approximation and projection (UMAP) algorithm (67) (Seurat implementation) followed by unsupervised graph-based clustering. Clusters of T cells were iteratively filtered based upon expression of *PTPRC* and *CD3E* and a lack of expression of myeloid markers, such as *CD14* and *SPI1*. Filtered cells were reanalyzed using the RPCA method followed by PCA for both the RNA and ADT assays. Finally, WNN was used to integrate the reanalyzed RNA and ADT assays followed by UMAP visualization. The final optimal number of PCs and clustering resolution were 30 and 0.9, respectively. T cell states were determined based upon select RNA and protein markers (Supplementary Table S2) and differentially expressed genes of each cluster determined using the Wilcoxon rank-sum test-based function. These annotations were supported by single cell gene enrichment analysis, which was performed as previously described using the escape R package (68–70). In brief, cluster specific gene signatures were derived from the top 50 differentially expressed genes from each cluster. Defined gene signatures from various single cell data sets of T cells were collected (17,33,69,71,72). Each cell was then assigned a score for each gene signature and the correlation of each cluster specific gene signature was assessed against other defined gene signatures. These correlations were used to support existing T cell annotations or were used to label specific T cell clusters (Supplementary Data 2).

The RNA assay from Cohort 1 was integrated with the RNA assay from Cohort 2, which was subjected to the same initial filtering parameters, using the RPCA method. Similarly, PCA was performed followed by UMAP visualization. In addition, T cells were iteratively filtered as previously specified. The final optimal number of PCs and clustering resolution were 40 and 1.3, respectively. T cell states were determined based upon select RNA markers, differentially expressed genes of each cluster, and guided by T cell annotations from Cohort 1, which were derived from both RNA and protein markers.

CD8^+^ T cells were subsetted based upon both RNA and protein expression (cohort 1) or just RNA expression of CD8 (cohort 1 & cohort 2). Samples were reintegrated with the RPCA method followed by UMAP visualization. The final optimal number of PCs and clustering resolution were 24 and 1.1, respectively (cohort 1) and 25 and 2.5, respectively (cohort 1 + cohort 2). T cell states were determined based upon select RNA markers, differentially expressed genes of each cluster, and guided by T cell annotations from the parent data set.

### Multiple cancer data set integration

CD8^+^ T cells from this current study of HGG and BrMet samples were integrated with CD8^+^ T cells from the following data sets: 1) a recent pan-cancer TIL data set (33), 2) a melanoma data set (34), 3) an independent GBM data set (17), and 4) a PDAC data set (35). Only samples with 5’ gene expression libraries (10X Genomics) were used for this integration. In several samples, genes in the Ensembl format were converted to respective gene names using clusterProfiler (73) and any outstanding discrepancies in gene names were corrected. Any genes related to T cell receptor genes, such as *TRAV* related genes, immunoglobulin genes, such as *IGH* related genes, and *MALAT1* were removed as previously described (33). Only genes common to all samples were used to avoid any potential library inconsistencies, with a final 10,211 genes shared by all samples. Prior to integration, T cells were separated by tumor type and the total number of T cells from each tumor type was randomly subsetted to 1,904 cells for a total of 39,984 cells. All samples were then merged, normalized, the top 2000 variable genes were calculated, and scaled. To minimize noise from cell cycle and tissue dissociation, the S phase score and G2M phase score calculated by the function *CellCycleScoring* and mitochondrial gene percentage calculated with *PercentageFeatureSet* were regressed (28). PCA was then performed with the optimal number of PCs followed by integration using the Harmony R package (74). Dimensionality reduction and visualization were performed with the uniform manifold approximation and projection (UMAP) algorithm (67) (Seurat implementation) followed by unsupervised graph-based clustering. The final optimal number of PCs and clustering resolution were 50 and 1.0, respectively. T cell states were determined based upon select RNA markers (Supplementary Table S2) and differentially expressed genes of each cluster determined using the Wilcoxon rank-sum test-based function (Supplementary Data S1: Sheet 5). These annotations were supported by the cell annotations specified in the meta data provided by the original authors.

### Exhaustion module score

The two exhaustion scores were generated using *AddModuleScore* (Seurat implementation) and based upon select gene markers and the gene list previously published by Tirosh *et al.* (28) (Supplementary Table S2).

### Pseudotemporal ordering analysis

Wang *et al.* HGG and BrMet data set:

Pseudotime trajectory analysis was performed with the monocle2 R package (31). Out of the CD8^+^ T cells, only the C2: CD8^+^ T_N_, C9: CD8^+^ *NR4A2*^lo^ *GZMK*^+^ T_Eff_, C10: CD8^+^ *NR4A2*^hi^ *GZMK*^+^ T_Eff_, C11: CD8^+^ cytotoxic T_RM_, and C12: CD8^+^ T_EMRA_ clusters were analyzed. The top 1,000 unique differentially expressed genes of these clusters were used to guide pseudotime trajectory analysis. The trajectory was rooted in the C2:CD8^+^ T_N_ cluster. The top 200 genes from the *differentialGeneTest* monocle2 function were used to generate the heatmap. Integrated pan cancer data set:

Pseudotemporal ordering analysis was performed with the monocle2 R package (31). Out of the CD8^+^ T cells, only the C1: T_N_, C2: *IL7R*^+^ T_EM_, C3: *NR4A2*^lo^ *GZMK*^+^ T_Eff_, C4: *NR4A2*^hi^ *GZMK*^+^ T_Eff_, C5: cytotoxic T_RM_, C7: T_EMRA_, and C8: T_Exhausted_ clusters were analyzed. The top 200 unique differentially expressed genes of these clusters were used to guide pseudotime trajectory analysis. The trajectory was rooted in the C1:CD8^+^ T_N_ cluster. The top 200 genes from the *differentialGeneTest* monocle2 function were used to generate the heatmap.

### Gene functional enrichment analysis

Gene functional enrichment analysis was performed through ToppGene (32). The top 25 DEGs expressed by CD8^+^ *GZMK*^+^ T cells were selected, and the top two unique gene ontology biological pathways were displayed. The pathways and associated genes are listed in Additional File 3: Sheet 2.

### Single Cell TCR analysis

Raw sequencing data generated from V(D)J libraries from both cohorts 1 and 2 were processed with the CellRanger V(D)J pipeline (10X Genomics, default settings, version 2.0.0) and mapped onto a human genome reference (GrCh38-2.0.0). The scRepertoire R package (75) was used for all subsequent clonotype analysis. Clonotype states were defined as the following: single (0 < X ≤ 1), doublet (1 < X ≤ 2), expanded (2 < X ≤ 50), and hyperexpanded (50 < X), where X is the number of cells in which the clonotype is detected.

### G-Rex Culturing & Bulk TCR Sequencing

Tumor samples were processed into 1 mm^3^ sections and incubated in a G-Rex 6M cell culture system (Wilson Wolf, Minnesota). Approximately 5-6 sections were incubated per well. Tumor samples were incubated in R10 media consisting of RPMI-1640 (Gibco) supplemented with 10% fetal bovine serum (FBS), 1% L-glutamine (Corning), 1% Pen/Strep (Gibco), 1% sodium pyruvate (Gibco), 0.5% sodium bicarbonate (Corning), 50 μM β-mercaptoethanol (Sigma-Aldrich), and 5,000 IU/mL recombinant human interleukin-2 (rhIL-2). Fresh IL-2 containing R10 media was added every 3 to 4 days. After approximately one month, cells were harvested by vigorous pipetting and then filtered through a 70 micron filter. Cells were then washed twice with R10 media and resuspended in ACK Lysis Buffer (Lonza Biosciences) if necessary. Cells were then frozen down in liquid nitrogen in 90% FBS with 10% DMSO.

Bulk T cell receptor (TCR) variable beta chain immunosequencing of genomic DNA from frozen T cells was performed using the ImmunoSEQ Assay (Adaptive Biotechnologies, Seattle, WA) on the following: TIL processed the day of surgical resection (D0), PBMCs isolated the day of surgical resection (D0), and expanded TIL (DX).

### Bulk TCR from G-rex Analysis

Data generated from the G-Rex systems were analyzed using the immunarch R package (76). Bulk TCRβ data was matched to single cell V(D)J data by matching bulk TCRβ sequences with single cell TCR clonotypes that contained the respective beta chains.

### Data availability

FASTQ files are available in the NCBI Sequence Read Archive (SRA) (SUB13030158). Processed gene expression and V(D)J matrices and Seurat objects used for all analyses are available on the open-access data sharing platform Zenodo (https://doi.org/10.5281/zenodo.8198492) (77).

### Statistical analysis

For comparison of groups, the Mann-Whitney U test was used. For comparison of frequencies, the Fisher’s exact test was used. For differential gene expression, the Wilcoxon rank-sum test (two-sided) was used with Bonferroni correction. p < 0.05 was considered to be statistically significant. Statistical analyses were performed with R statistical language V 4.1.0 or Seurat.

## Acknowledgements

We thank the Genome Technology Access Center at the McDonnell Genome Institute at Washington University School of Medicine for help with genomics services. The Center is partially supported by NCI Cancer Center Support Grant #P30 CA91842 to the Siteman Cancer Center and by ICTS/CTSA Grant# UL1TR002345 from the National Center for Research Resources (NCRR), a component of the National Institutes of Health (NIH), and NIH Roadmap for Medical Research. This publication is solely the responsibility of the authors and does not necessarily represent the official view of NCRR or NIH.

AZW was supported by the Medical Scientist Training Program at Washington University in St. Louis.

## Funding

Cancer Research Institute Lloyd J. Old STAR Award (GPD)

Paul Calabresi K12 Career Development Award for Clinical Oncology (TMJ)

## Author contributions

Conceptualization: MOS, TMJ, GPD

Formal analysis: AZW, BLM, SMK, MAC

Funding acquisition: TMJ, GPD

Investigation: AZW, BLM, MOS, LAL, AJL, JDB, ECL, AHK, JLD, MRC, PSJ, BDC, DPC, BSC

Methodology: MOS, AJL, AZW, SMK

Project administration: AZW, GPD, TMJ, AAP

Resources: AZW, MOS, LAL, AJL, JDB, AAP, TMJ, GPD

Software: AZW, BLM, MAC, SMK, AAP

Supervision: GPD, TMJ, AAP

Visualization: AZW, NDS, ML

Writing – original draft: AZW

Writing – review & editing: AZW, GPD, TMJ, AAP

## Notes

**Conflict of interest statement** DPC has received financial compensation from the Massachusetts Institute of Technology, Advise Connect Inspire, German Accelerator, Lilly, GlaxoSmithKline, Incephalo, Boston Pharmaceuticals, Boston Scientific and Pyramid Biosciences (equity interest) for advisory input. He has also received financial compensation and travel reimbursement from Merck for invited lectures, and from the US NIH and DOD for clinical trial and grant review. The remaining authors declare that they have no competing interests.

### Competing Interest Statement

DPC has received financial compensation from the Massachusetts Institute of Technology, Advise Connect Inspire, German Accelerator, Lilly, GlaxoSmithKline, Incephalo, Boston Pharmaceuticals, Boston Scientific and Pyramid Biosciences (equity interest) for advisory input. He has also received financial compensation and travel reimbursement from Merck for invited lectures, and from the US NIH and DOD for clinical trial and grant review.
The remaining authors declare that they have no competing interests.

