## Supplementary Figures and Tables for "Glioblastoma-infiltrating CD8^+^ T cells are predominantly a clonally expanded *GZMK*^+^ effector population"

| **Patient ID** | **Sex** | **Age** | **Pathology** | **MGMT Status** | **TIL** | **PBMC** | **G-Rex** | **Cohort** | **Analysis** |
| --- | --- | --- | --- | --- | --- | --- | --- | --- | --- |
| **GBM091** | **F** | **68** | **IDH WT Primary GBM** | **Indeterminate** | **Y** | **N** | **N** | **1** | **5’ CITE** |
| GBM098 | F | 59 | IDH WT Primary GBM | Unmethylated | Y | N | N | 1 | 5’ CITE, V(D)J |
| **GBM104Re*** | **M** | **33** | **IDH WT Recurrent GBM** | **Unmethylated** | **Y** | **N** | **N** | **1** | **5’ CITE, V(D)J** |
| GBM105 | M | 64 | IDH WT Primary GBM | Methylated | Y | N | Y | 1 | 5’ CITE, V(D)J, bulk TCR |
| **GBM106** | **M** | **24** | **IDH WT Primary GBM** | **Methylated** | **Y** | **Y** | **Y** | **1** | **5’ CITE, V(D)J, bulk TCR** |
| GBM111Re* | M | 33 | IDH WT Recurrent GBM | Unmethylated | Y | Y | Y | 1 | 5’ CITE, V(D)J, bulk TCR |
| **GBM113** | **F** | **65** | **IDH WT Primary GBM** | **Methylated** | **Y** | **Y** | **Y** | **1** | **5’ CITE, V(D)J, bulk TCR** |
| GBM114 | F | 72 | IDH WT Primary GBM | Unmethylated | Y | Y | Y | 1 | 5’ CITE, V(D)J, bulk TCR |
| **GBM115** | **M** | **56** | **IDH WT Primary GBM** | **Unmethylated** | **Y** | **N** | **N** | **1** | **5’ CITE, V(D)J** |
| G4A112Re | M | 44 | IDH Mut  Recurrent Grade 4 Astrocytoma | Methylated | Y | Y | Y | 1 | 5’ CITE, V(D)J, bulk TCR |
| **GBM056** | **F** | **55** | **IDH WT Primary GBM** | **Unmethylated** | **Y** | **N** | **N** | **2** | **5’ GEX, V(D)J** |
| GBM063 | M | 55 | IDH WT Primary GBM | Unmethylated | Y | N | N | 2 | 5’ GEX, V(D)J |
| **GBM064** | **F** | **55** | **IDH WT Primary GBM** | **Unmethylated** | **Y** | **N** | **N** | **2** | **5’ GEX, V(D)J** |
| GBM074 | M | 51 | IDH WT Primary GBM | Unmethylated | Y | N | N | 2 | 5’ GEX, V(D)J |
| **G4A062** | **F** | **49** | **IDH Mut**  **Primary Grade 4 Astrocytoma** | **Methylated** | **Y** | **N** | **N** | **2** | **5’ GEX, V(D)J** |
| G4A065Re | M | 25 | IDH Mut  Recurrent Grade 4 Astrocytoma | Indeterminate | Y | N | N | 2 | 5’ GEX, V(D)J |
| **BrMet009** | **M** | **58** | **NSCLC** | **N/A** | **Y** | **N** | **N** | **2** | **5’ GEX, V(D)J** |
| BrMet010 | F | 57 | NSCLC | N/A | Y | N | N | 2 | 5’ GEX, V(D)J |
| **BrMet018** | **F** | **59** | **Breast** | **N/A** | **Y** | **N** | **N** | **2** | **5’ GEX, V(D)J** |
| BrMet027 | F | 31 | Breast | N/A | Y | N | N | 2 | 5’ GEX, V(D)J |
| **BrMet028** | **M** | **69** | **Melanoma** | **N/A** | **Y** | **N** | **N** | **2** | **5’ GEX, V(D)J** |
| **MGH_GBM001** | **F** | **49** | **IDH WT Primary GBM** | **Unmethylated** | **N** | **Y** | **N** | **N/A** | **Flow cytometry** |
| MGH_GBM004 | M | 91 | IDH WT Primary GBM | Methylated | N | Y | N | N/A | Flow cytometry |
| **MGH_GBM012** | **F** | **66** | **IDH WT Primary GBM** | **Unmethylated** | **Y** | **Y** | **N** | **N/A** | **Flow cytometry** |
| MGH_GBM013 | F | 81 | IDH WT Primary GBM | Methylated | N | Y | N | N/A | Flow cytometry |
| **MGH_GBM014** | **F** | **82** | **IDH WT Primary GBM** | **Unmethylated** | **Y** | **Y** | **N** | **N/A** | **Flow cytometry** |
| MGH_GBM015 | F | 64 | IDH WT Primary GBM | Methylated | Y | N | N | N/A | Flow cytometry |

**Table S1: Patient demographics and sample characteristics.** Clinical and sample details for all patient samples. WT: wild-type, Mut: mutant, NSCLC: non-small cell lung cancer, MGMT: O-6-methylguanine-DNA methyltransferase, 5’ CITE: 5’ Cellular indexing of transcriptomes and epitope single cell sequencing, 5’ GEX: 10X Genomics 5’ Gene expression single cell sequencing, V(D)J: 10X Genomics V(D)J single cell sequencing, * indicates same patient.

| **T cell state** | **Markers** |
| --- | --- |
| Naïve T cell (T_N_) | RNA: *CCR7, SELL, TCF7, IL7R-*  ADT: CD45RA |
| Central memory T cell (T_CM_) | RNA: *CCR7, SELL, TCF7, IL7R-*  ADT: CD45RO |
| Effector memory T cell (T_EM_) | RNA: *CCR7-, SELL-, TCF7-, IL7R*  ADT: CD69, CD45RO |
| Resident memory T cell (T_RM_) | RNA: *NR4A2, ITGAE*  ADT: CD103 |
| Effector memory RA T cell (T_EMRA_) | RNA: *CCR7-, SELL-, TCF7-, IL7R*  ADT: CD45RA |
| Regulatory T cell (T_reg_) | RNA: *FOXP3, CTLA4, TIGIT* |
| Proliferating T cell (T_proliferating_) | RNA: *MKI67, UBE2C, BIRC5, TOP2A* |
| Stress signature | *HSP* related genes |
| MAIT-like T cell | V(D)J: TRAV1-2.TRAJ33 TCRα chain |
| *GZMK*^+^ T_Eff_ | RNA: *GZMK, PRF1-, GZMB-, GNLY-, NR4A2*^lo/hi^  ADT: CD45RO, PD1, CD27, CD28  T_EM_ markers |
| Cytotoxic T_RM_ | RNA: *GZMB, GNLY, PRF1, IFNG, NKG7,*  T_RM_ markers |
| Natural killer-like (NK-like) | *FCER1G, NCAM1, KLRC2, NCR1* |
| IFNG stimulated T cells | *IFIT1, IFIT3, IFI6, MX1, ISG15* |
| RNA Inhibitory Score (citation) | *PDCD1, CTLA4, TIGIT, LAG3, HAVCR2, TNFSRF9, BTLA* |
| Tirosh Exhaustion Score (citation) | *CXCL13, TNFRSF1B, RGS2, TIGIT, CD27, TNFRSF9, SLA, RNF19A, INPP5F, XCL2, HLA-DMA, FAM3C, UQCRC1, WARS, EIF3L, KCNK5, TMBIM6, CD200, ZC3H7A, SH2D1A, ATP1B3, MYO7A, THADA, PARK7, EGR2, FDFT1, CRTAM, IFI16* |

**Table S2. Gene markers for cell type identification and gene signatures.**

| **Antibody** | **Source** | **Clone #** | **Barcode** |
| --- | --- | --- | --- |
| CD45RO, PerCP-Cy5.5 | Biolegend | UCHL1 | N/A |
| CD45, PerCP-Cy5.5 | Biolegend | HI30 | N/A |
| CD3, PE/Cy7 | Biolegend | UCHT1 | N/A |
| CD3, APC | Biolegend | UCHT1 | N/A |
| CD3, FITC | Biolegend | UCHT1 | N/A |
| CD4, BV605 | Biolegend | OKT4 | N/A |
| CD8, BV650 | Biolegend | SK1 | N/A |
| CD8, PE/Cy7 | Biolegend | SK1 | N/A |
| CD11b, PE | Biolegend | ICRF44 | N/A |
| TCR Vα7.2, BV711 | Biolegend | 3C10 | N/A |
| TCR Vα7.2, Alexa Fluor 700 | Biolegend | 3C10 | N/A |
| GZMK, PE | Biolegend | GM26E7 | N/A |
| GZMB, FITC | Biolegend | QA16A02 | N/A |
| Zombie Aqua | Biolegend | N/A | N/A |
| Zombie NIR | Biolegend | N/A | N/A |
| CD274 (B7-H1, PD-L1) | Biolegend | 29E.2A3 | GTTGTCCGACAATAC |
| CD270 (HVEM, TR2) | Biolegend | 122 | TGATAGAAACAGACC |
| CD70 | Biolegend | 113-16 | CGCGAACATAAGAAG |
| CD30 | Biolegend | BY88 | TCAGGGTGTGCTGTA |
| CD154 | Biolegend | 24-31 | GCTAGATAGATGCAA |
| CD4 | Biolegend | SK3 | GAGGTTAGTGATGGA |
| CD8 | Biolegend | SK1 | GCGCAACTTGATGAT |
| CD56 (NCAM) | Biolegend | 5.1H11 | TCCTTTCCTGATAGG |
| CD45RA | Biolegend | HI100 | TCAATCCTTCCGCTT |
| CD25 | Biolegend | BC96 | TTTGTCCTGTACGCC |
| CD45RO | Biolegend | UCHL1 | CTCCGAATCATGTTG |
| CD279 (PD-1) | Biolegend | EH12.2H7 | ACAGCGCCGTATTTA |
| TIGIT (VSTM3) | Biolegend | A15153G | TTGCTTACCGCCAGA |
| CD103 (Integrin αE) | Biolegend | Ber-ACT8 | GACCTCATTGTGAAT |
| CD69 | Biolegend | FN50 | GTCTCTTGGCTTAAA |
| CD62L | Biolegend | DREG-56 | GTCCCTGCAACTTGA |
| CD152 (CTLA-4) | Biolegend | BNI3 | ATGGTTCACGTAATC |
| CD223 (LAG-3) | Biolegend | 11C3C65 | CATTTGTCTGCCGGT |
| KLRG1 (MAFA) | Biolegend | 2F1/KLRG1 | GTAGTAGGCTAGACC |
| CD27 | Biolegend | O323 | GCACTCCTGCATGTA |
| CD107a (LAMP-1) | Biolegend | H4A3 | CAGCCCACTGCAATA |
| CD95 (Fas) | Biolegend | DX2 | CCAGCTCATTAGAGC |
| CD134 (OX40) | Biolegend | Ber-ACT35 (ACT35) | AACCCACCGTTGTTA |
| HLA-DR | Biolegend | L243 | AATAGCGAGCAAGTA |
| CD57 Recombinant | Biolegend | QA17A04 | AACTCCCTATGGAGG |
| CD366 (Tim-3) | Biolegend | F38-2E2 | TGTCCTACCCAACTT |
| CD272 (BTLA) | Biolegend | MIH26 | GTTATTGGACTAAGG |
| CD278 (ICOS) | Biolegend | C398.4A | CGCGCACCCATTAAA |
| CD96 (TACTILE) | Biolegend | NK92.39 | TGGCCTATAAATGGT |
| CD39 | Biolegend | A1 | TTACCTGGTATCCGT |
| CD244 (2B4) | Biolegend | C1.7 | TCGCTTGGATGGTAG |
| CD137 (4-1BB) | Biolegend | 4B4-1 | CAGTAAGTTCGGGAC |
| CD357 (GITR) | Biolegend | 108-17 | ACCTTTCGACACTCG |
| CD48 | Biolegend | BJ40 | CTACGACGTAGAAGA |
| CD161 | Biolegend | HP-3G10 | GTACGCAGTCCTTCT |
| CD226 (DNAM-1) | Biolegend | 11A8 | TCTCAGTGTTTGTGG |
| CD49b | Biolegend | P1E6-C5 | GCTTTCTTCAGTATG |
| CD28 | Biolegend | CD28.2 | TGAGAACGACCCTAA |
| CD38 | Biolegend | HIT2 | TGTACCCGCTTGTGA |
| CD127 (IL-7Rα) | Biolegend | A019D5 | GTGTGTTGTCCTATG |
| B7-H4 | Biolegend | MIH43 | TGTATGTCTGCCTTG |
| CD36 | Biolegend | 5-271 | TTCTTTGCCTTGCCA |
| CD73 (Ecto-5'-nucleotidase) | Biolegend | AD2 | CAGTTCCTCAGTTCG |

**Table S3. Antibody markers and respective clone numbers.**

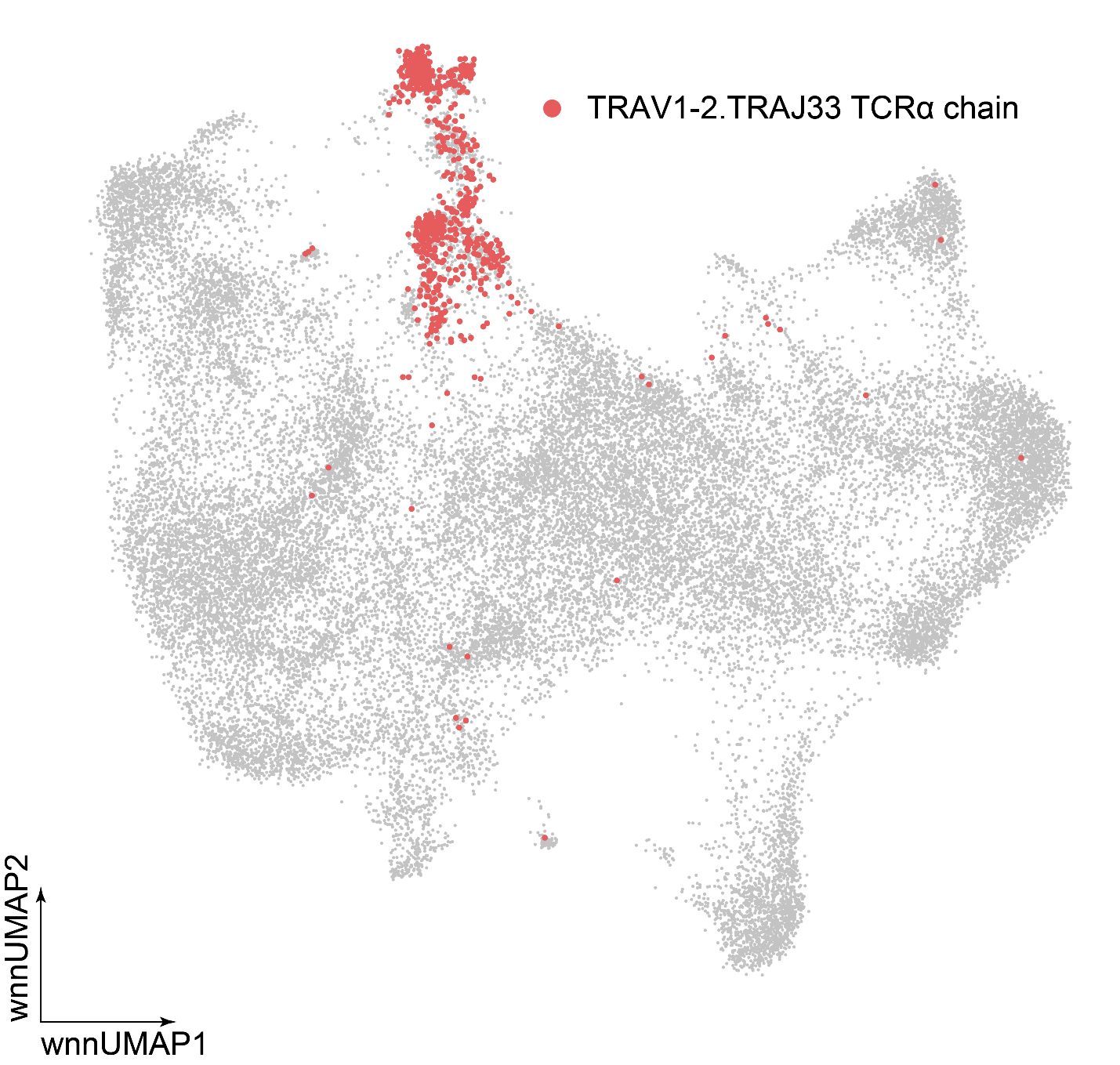

**Fig. S1. MAIT clonotype occurrence in T cells from cohort 1.** wnnUMAP visualization of the TRAV1-2.TRAJ33 TCRα chain, the alpha chain of the semi-invariant αβ TCR of MAIT cells.

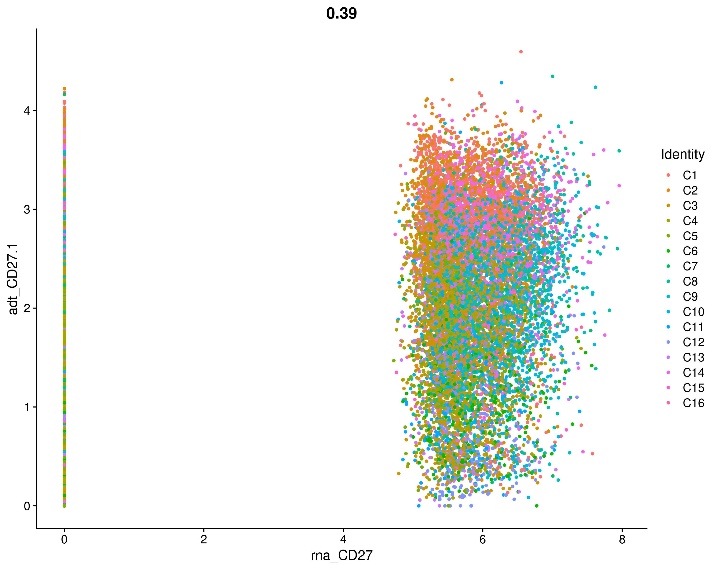

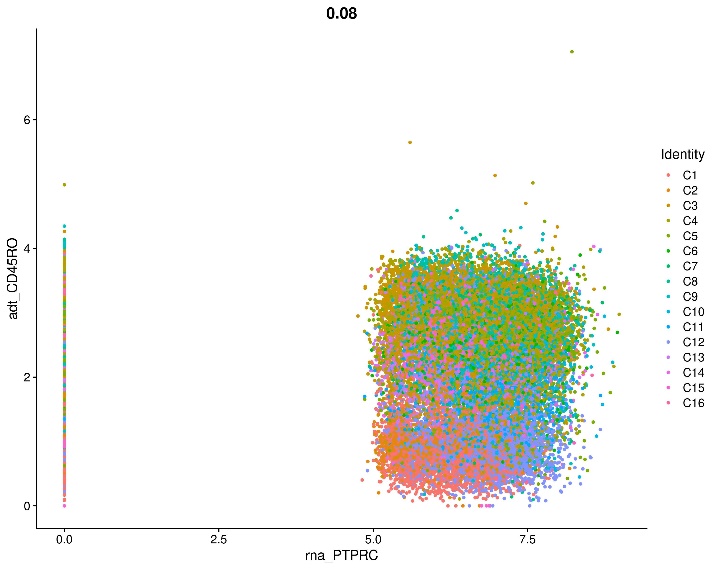

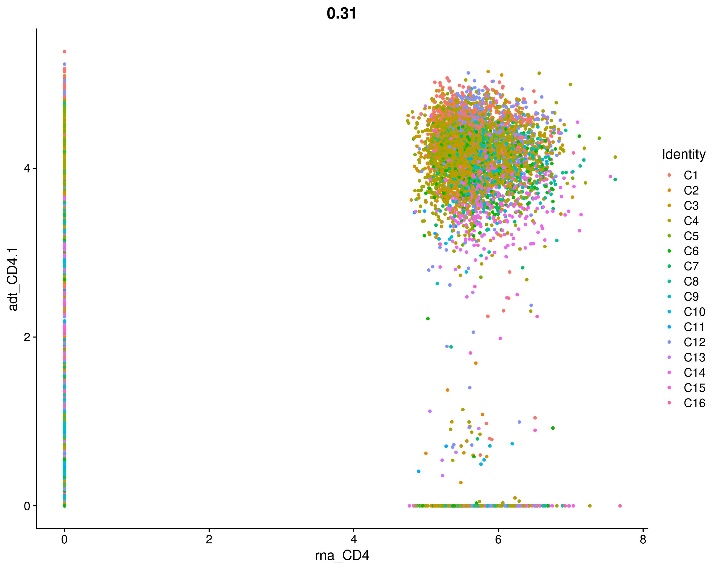

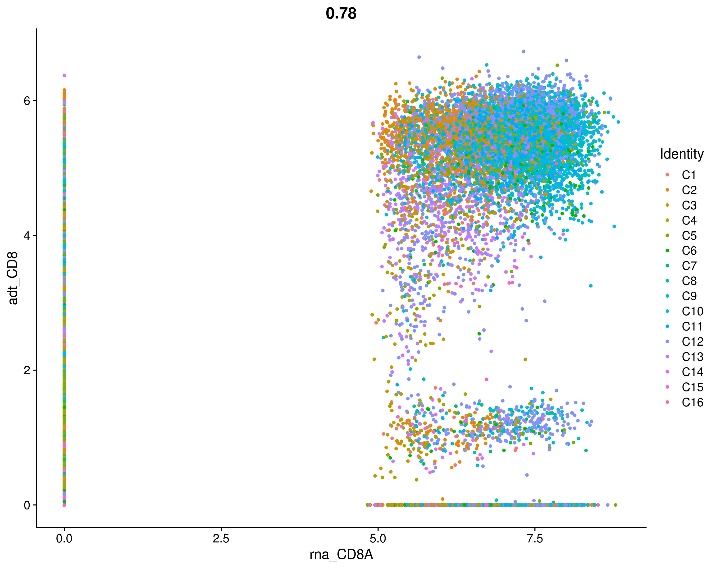

**Fig. S2. Correlation plots of expression of various proteins and respective genes by T cells from cohort 1.** Correlation plots of single cell protein (ADT) vs. RNA expression for several genes/proteins from cohort 1 patient samples.

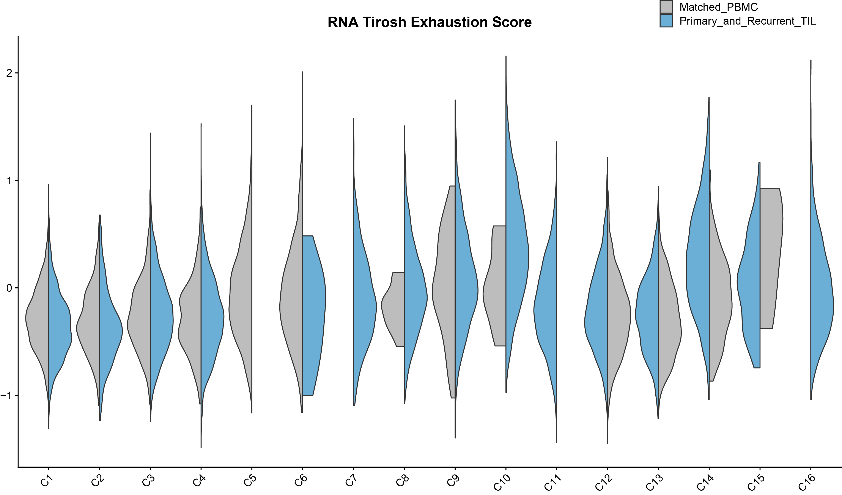

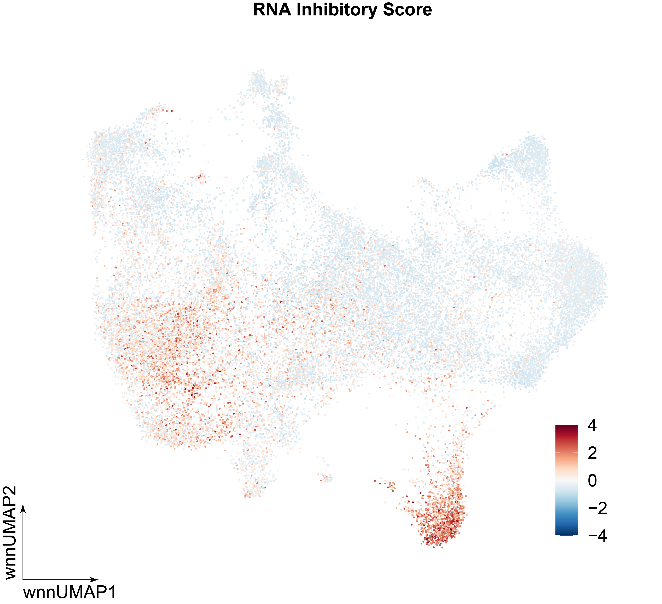

**B**

**A**

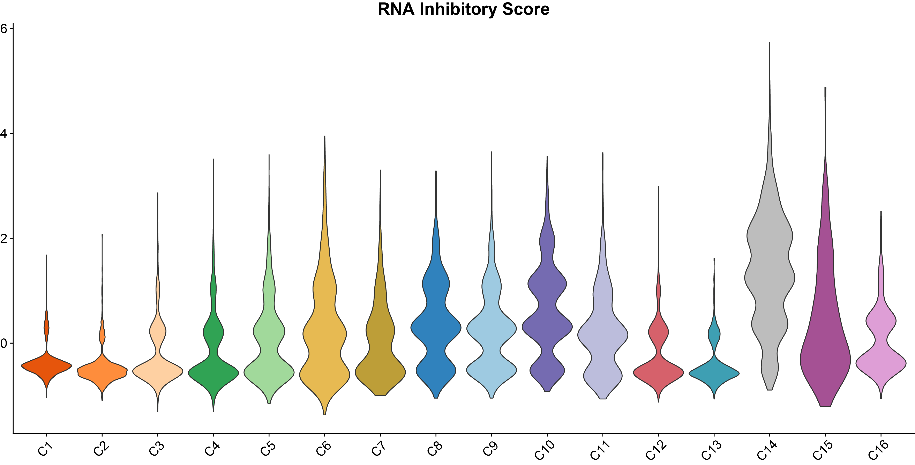

**C**

**D**

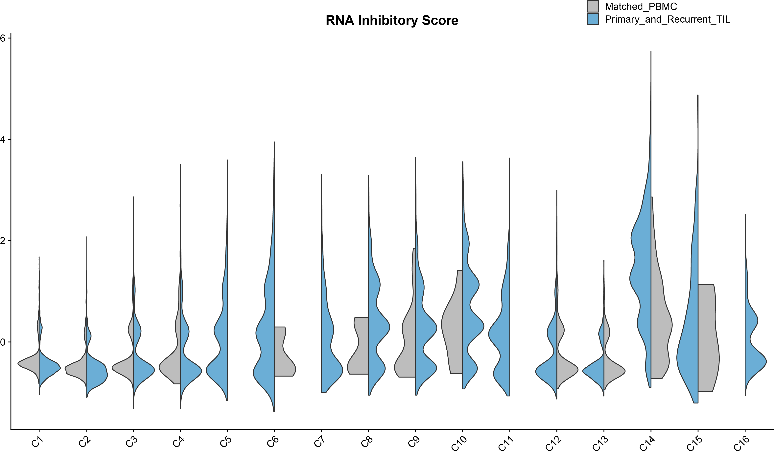

**Fig. S3. Analysis of RNA Tirosh exhaustion and inhibitory scores expressed by T cells from cohort 1. (A)** Violin plot of RNA Tirosh exhaustion score expression by each T cell state split by sample type. **(B)** UMAP visualization of RNA inhibitory score expression. **(C)** Violin plot of RNA inhibitory score expression with each T cell state as colored in Fig. 1B. **(D)** Violin plot of RNA inhibitory score expression by each T cell state split by sample type.

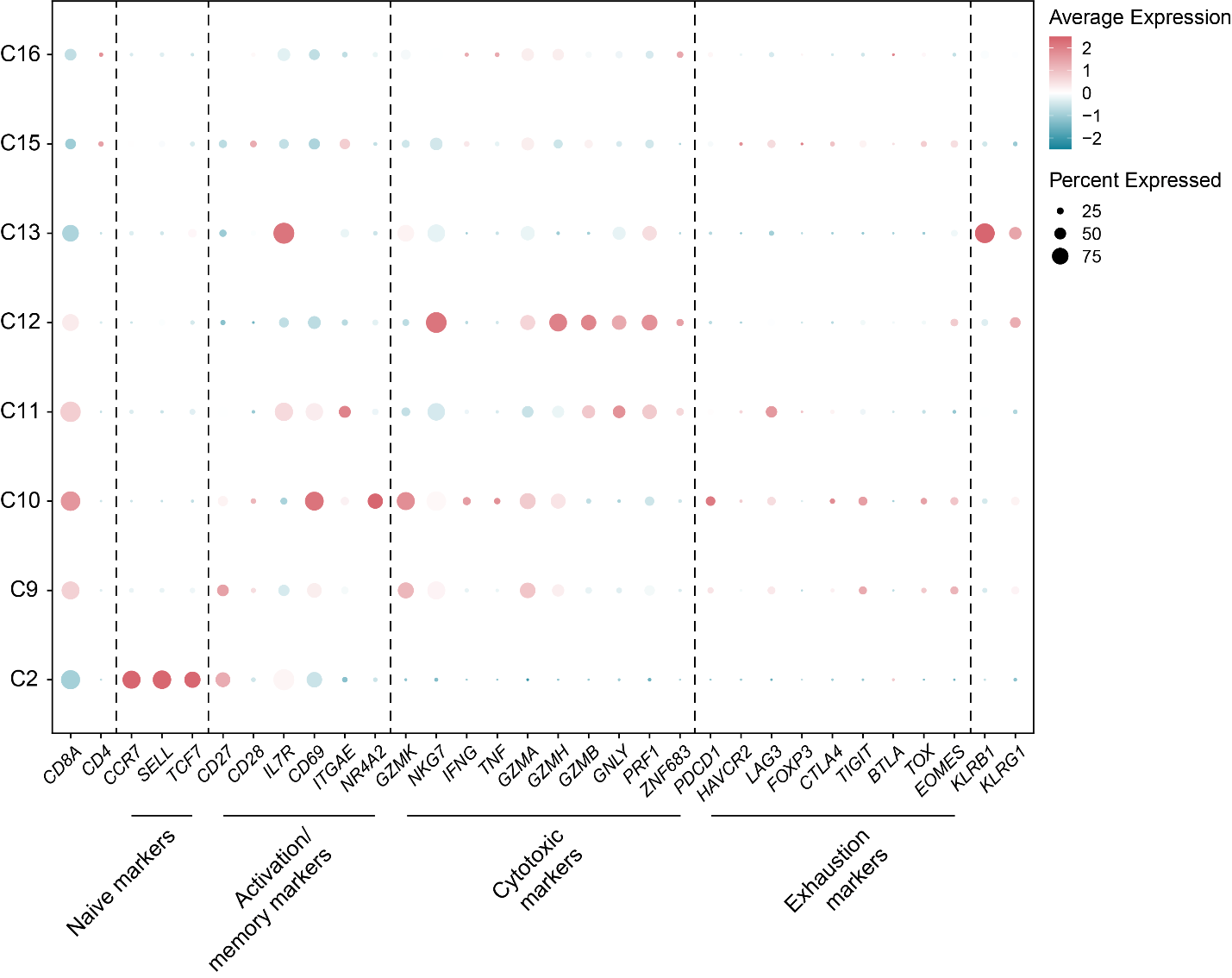
**Fig. S4. Dot plot of select gene markers for cell type identification of CD8^+^ T cells from cohort 1.** Dot plot of expression of select gene markers used to identify CD8^+^ T cell subsets as shown in Fig. 2A.

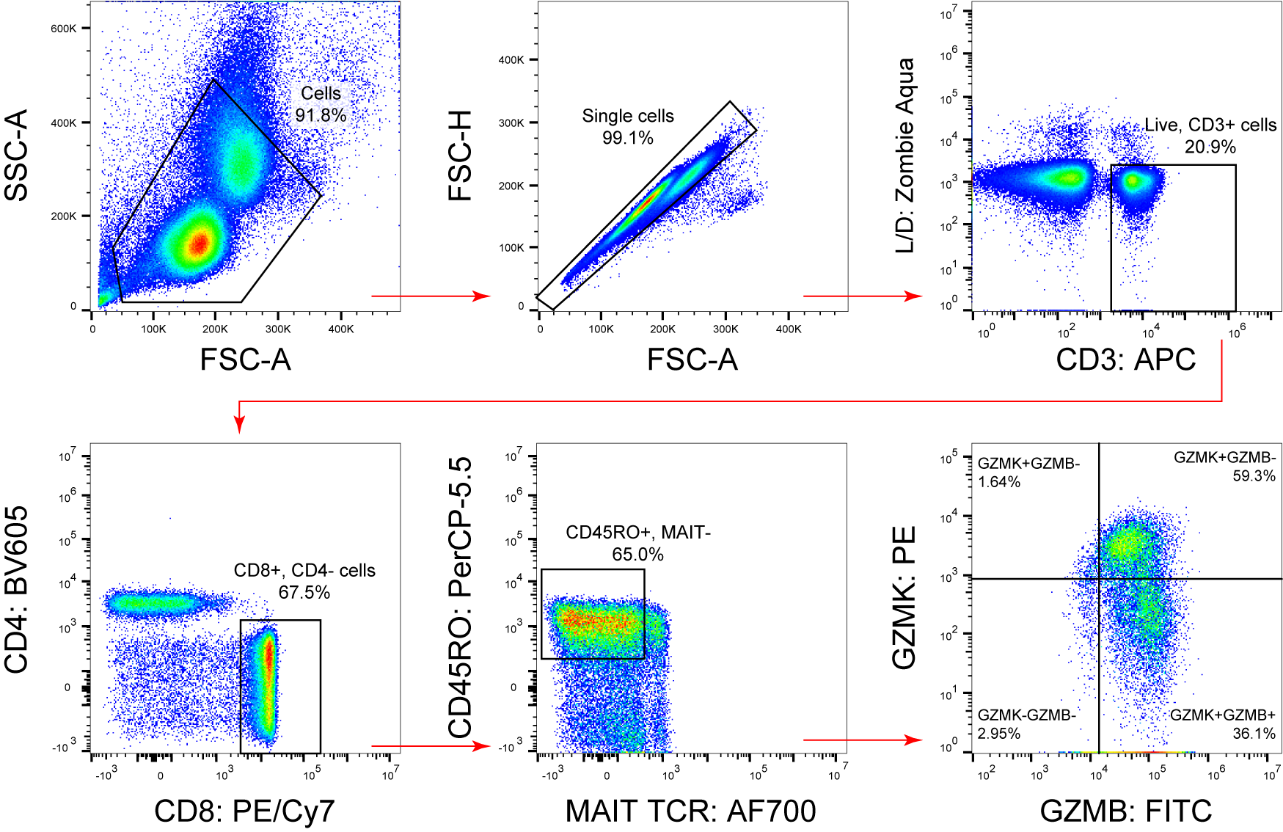

**Fig. S5. Gating strategy of GBM and PBMC T cells.** Gating strategy used to determine frequency of GZMK and GZMB populations.

**A**

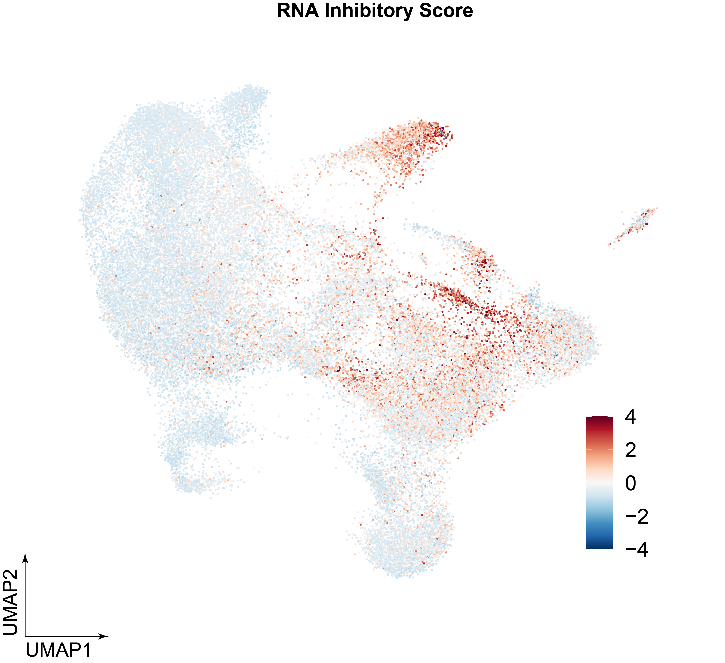

**B**

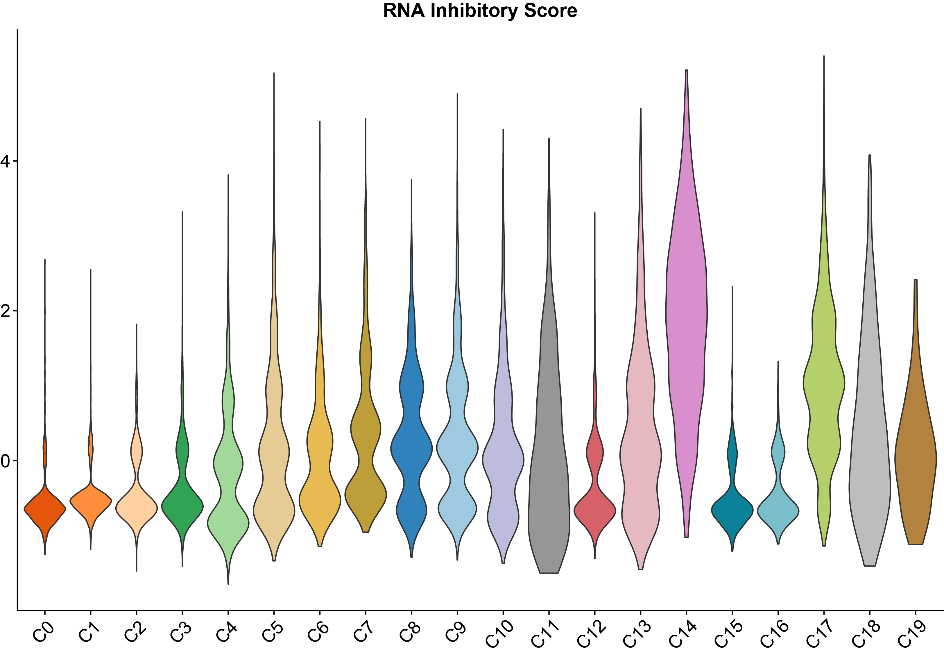

**Fig. S6. Inhibitory score analysis of the integrated T cells from cohorts 1 and 2. (A)** UMAP visualization of RNA inhibitory score expression. **(B)** Violin plot of RNA inhibitory score expression with each T cell state as colored in Fig. 3A.

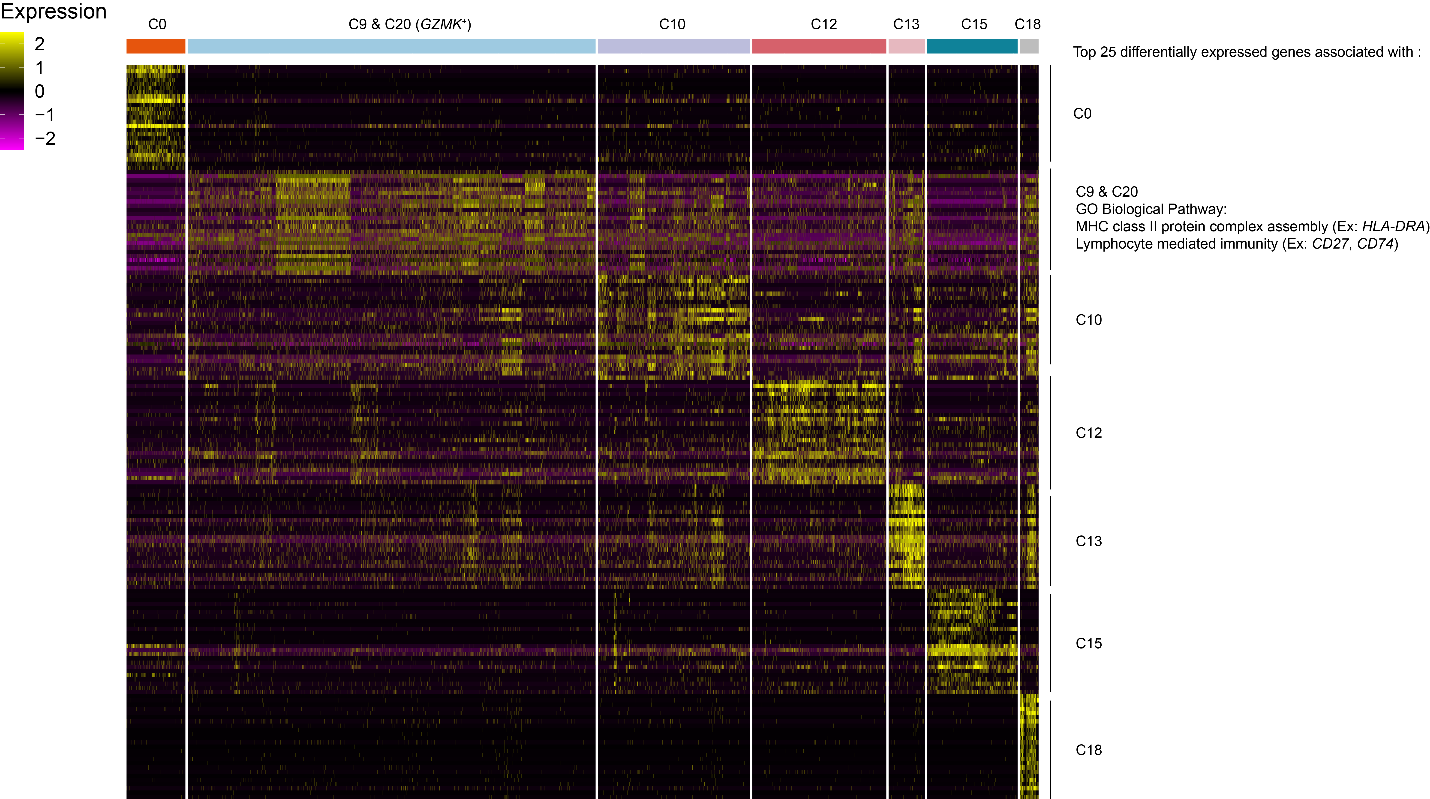

**Fig. S7. Heatmap of top differentially expressed genes by CD8^+^ T cell states from cohorts 1 and 2.** Heatmap of the top 25 differentially expressed genes expressed by CD8^+^ T cell clusters from Fig. 3F. The top 25 differentially expressed genes of *NR4A2*^lo/hi^ *GZMK*^+^ T cells were used for gene enrichment analysis. Two of the top associated biological pathways included “MHC class II protein complex assembly” and “lymphocyte mediated immunity”.

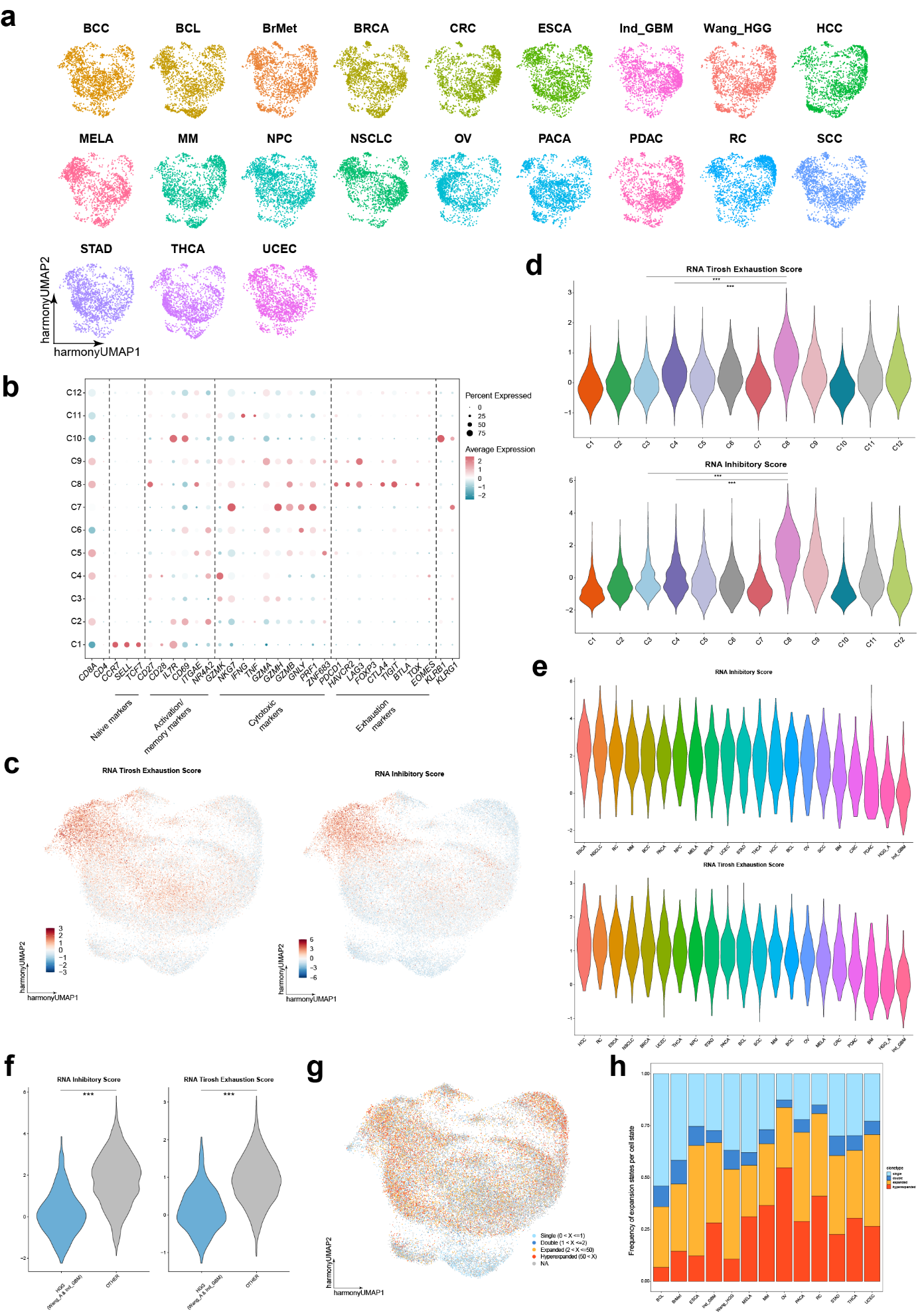

**Fig. S8. Harmony integration of TIL from many cancer types. (A)** Harmony UMAP visualization with different cancer types highlighted. **(B)** Dot plot of expression of select gene markers to identify CD8^+^ T cell subsets as shown in Fig. 4A. **(C)** UMAP visualizations of RNA inhibitory and Tirosh exhaustion score expression. **(D)** Violin plots of RNA inhibitory and Tirosh exhaustion score expression with each T cell state as colored in Fig. 4A. **(E)** Violin plots of RNA inhibitory and Tirosh exhaustion score expression by exhausted T cells (C8) with tumor type colored. **(F)** Violin plots comparing the RNA inhibitory and Tirosh exhaustion scores between exhausted T cells (C8) derived from HGG (both Ind_GBM and current HGG dataset) and all other tumor types. **(G)** Harmony UMAP visualization with clonotype enrichment states highlighted. **(H)** Stacked bar plot visualizing relative frequency of each clonotype enrichment state per cancer type. In **(F)**, significance calculated using Mann-Whitney U-test. ***p<2.2e^-16^. Ind_GBM = independent GBM data set.

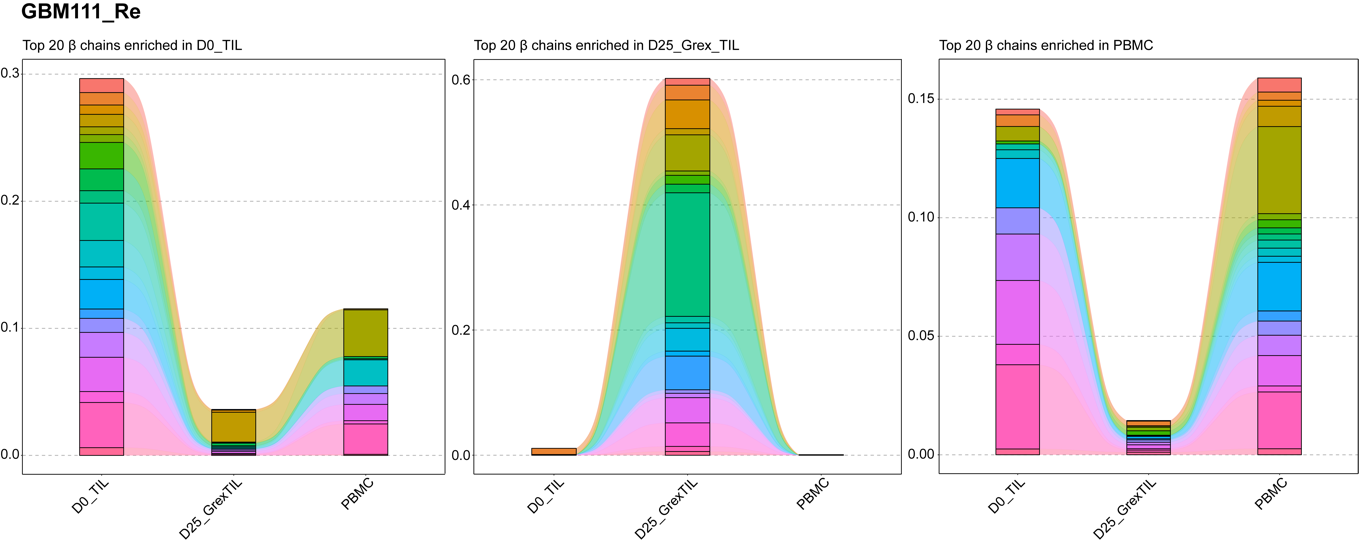

**B**

**A**

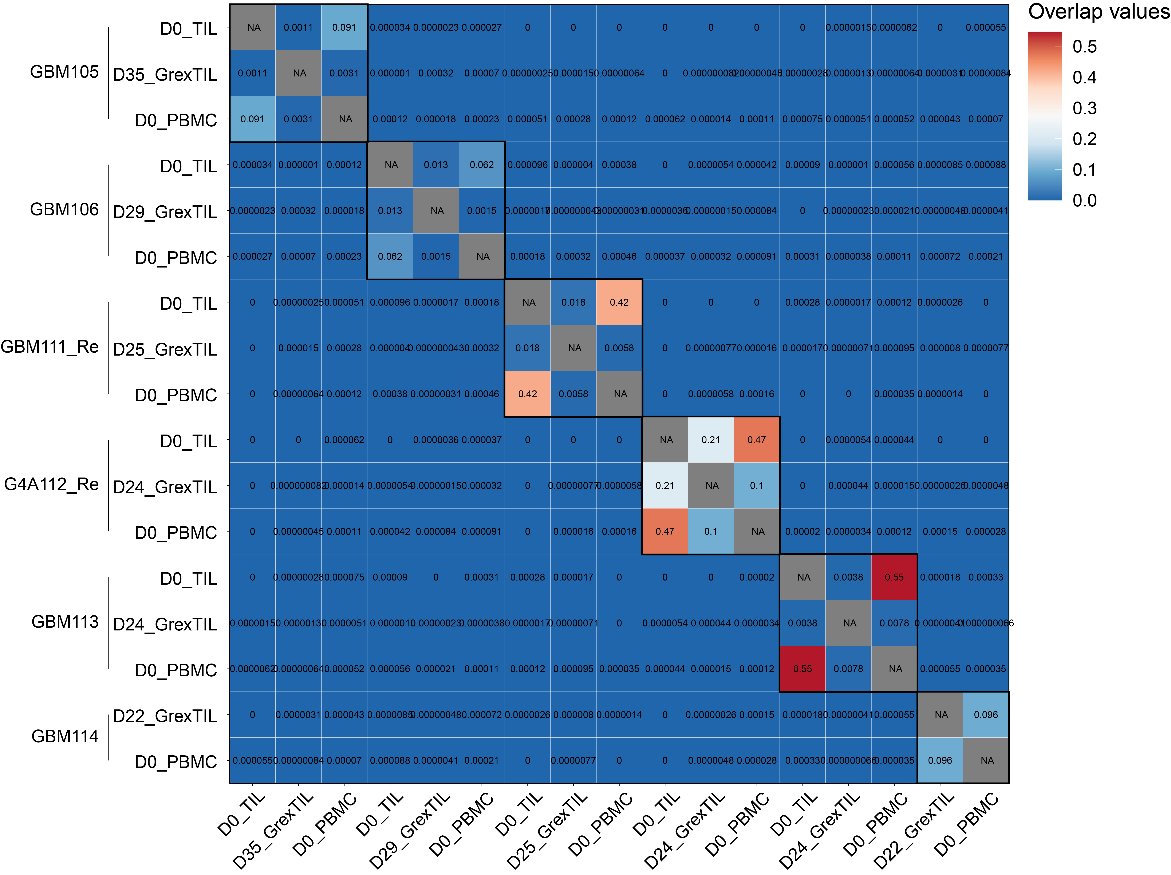

**Fig. S9. TCR landscape of *ex vivo* expanded TIL does not recapitulate the initial TCR landscape. (A)** Alluvial plots highlighting the overlap among matched sample types of the top 20 β chains in D0_TIL, DX_G-Rex_TIL, and PBMC, respectively. **(B)** Morisita’s index overlap of β chain landscapes of all patient samples and sample types.
